# Mechanics of DNA Replication and Transcription Guide the Asymmetric Distribution of RNAPol2 and Nucleosomes on Replicated Daughter Genomes

**DOI:** 10.1101/553669

**Authors:** Rahima Ziane, Alain Camasses, Marta Radman-Livaja

## Abstract

Eukaryotic DNA replication occurs in the context of chromatin. Chromatin in its capacity as a transcription regulator, is also thought to have a role in the epigenetic transmission of transcription states from one cell generation to the next. It is still unclear how chromatin structure survives the disruptions of nucleosomal architecture during genomic replication or if chromatin features are indeed instructive of the transcription state of the underlying gene. We have therefore developed a method for measuring chromatin structure dynamics after replication – ChIP -NChAP (Chromatin Immuno-Precipitation - Nascent Chromatin Avidin Pulldown) - which we used to monitor the distribution of RNAPol2 and new and old H3 histones on newly-replicated daughter genomes in S. Cerevisiae. The strand specificity of our sequencing libraries allowed us to uncover the inherently asymmetric distribution of RNAPol2, H3K56ac (a mark of new histones), and H3K4me^3^ and H3K36me^3^ (“active transcription marks” used as proxies for old histones) on daughter chromatids. We find a difference in the timing of lagging and leading strand replication on the order of minutes at a majority of yeast genes. Nucleosomes and RNAPol2 preferentially bind to either the leading or the lagging strand gene copy depending on which one replicated first and RNAPol2 then shifts to the sister copy after its synthesis has completed. Our results suggest that active transcription states are inherited simultaneously and independently of their underlying active chromatin states through the recycling of the transcription machinery and old histones, respectively. We find that “active” histone marks do not instruct the cell to reestablish the same active transcription state at its underlying genes. We propose that rather than being a consequence of chromatin state inheritance transcription actually contributes to the reestablishment of chromatin states on both replicated gene copies. Our findings are consistent with a two-step model of chromatin assembly and RNAPol2 binding to nascent DNA that is based on local differences in replication timing between the lagging and leading strand. The model describes how chromatin and transcription states are first restored on one and then the other replicated gene copy, thus ensuring that after division each cell will have “inherited” a gene copy with identical gene expression and chromatin states.

## Introduction

All eukaryotic genomic processes happen in the context of chromatin. The smallest repeating subunit of chromatin is the nucleosome: a 147 bp DNA segment wrapped 1.65 turns around a histone octamer core, consisting of one H3/H4 tetramer and two H2A/H2B dimers(*1*). Since architectural features of chromatin limit the accessibility of the DNA substrate to DNA processing enzymes involved in replication, transcription or repair, chromatin -in addition to being a genome packaging system- is the foremost regulatory system for all DNA based processes. Regulatory mechanisms embedded in specific chromatin configurations include: nucleosome positioning along DNA, posttranslational histone modifications, histone variants, and higher order structures such as chromatin loops and topologically associated domains (for reviews see (*2–9*)).

Chromatin configuration is transiently disrupted with every round of genome replication. Nucleosomes bound to DNA, which will be referred to as old nucleosomes from here on, are disassembled ahead of the replication fork and recycled behind it on the two newly formed daughter chromatids (*10*). Concomitantly with old nucleosome recycling, new histones are assembled on daughter chromatids to restore optimal nucleosome density after genome duplication (for review see (*11*)). The process of nucleosome reassembly on daughter chromatids and the (re)establishment of a specific chromatin architecture on replicated gene copies is called chromatin maturation. In theory chromatin maturation should result in the accurate inheritance of a specific chromatin feature, which should then help preserve the expression state of the underlying gene for the next cell generation. Following the same logic, if the chromatin state of a gene is modified during genome replication, the transcription state of that gene should also be changed. This kind of epigenetic transformation is thought to be at the heart of cellular differentiation. Whether chromatin structure is indeed accurately inherited and whether it is instructive of the underlying gene expression state are however still open questions.

The cell is faced with two problems after every replication event: 1. it has to either restore its chromatin configuration to its pre-replication state on both daughter genomes in order to maintain the same transcription program or use the disruption caused by replication as an “opportunity” to modulate chromatin configuration (symmetrically on both genomes or asymmetrically only on one replicated copy) and change the transcription program in response to developmental or environmental signals; 2. it has to globally regulate transcription levels in response to gene copy number doubling after genome replication in G2/M, until cellular division restores the original gene copy number.

While it is largely accepted that old and new histones bind to both daughter chromatids after replication (*12, 13*), the precise distribution pattern is still not clear. Old and new histones could bind completely symmetrically and randomly to either chromatid or in locally asymmetrical segments, thus forming contiguous alternating “patches” of old and new nucleosomes on the same chromatid. The mode of old and new histone distribution has implications for the mechanism of restauration of chromatin states. A map of old nucleosome distribution on replicated chromatids would thus provide clues on how and if potential epigenetic information “embedded” in old nucleosomes might be copied to new nucleosomes. It would also shed light on how replicated gene copies might end up in different gene expression states, which presumably occurs during differentiation. Most importantly, it is still not known whether and how the distribution pattern of old and new histones affects transcription after replication. While recent studies have observed asymmetrical binding of old and newly synthesized histones to replicated daughter chromatids, (*14–16*), it is still not clear how RNAPol2 binding and consequently transcription of replicated gene copies might be affected by such asymmetrical chromatin assembly.

We have recently developed a high throughput sequencing based technique to map chromatin features on newly replicated DNA – Nascent Chromatin Avidin Pulldown (NChAP) (*17*). Using NChAP, we determined that nucleosome positioning maturation after replication depends on transcription. We also measured faster nucleosome repositioning on the copy replicated by the leading strand when gene transcription travels in the same direction as the replication fork (“same” orientation genes). On the other hand, nucleosomes repositioning is faster on the lagging strand copy for “opposite” orientation genes. This correlation between nucleosome repositioning rates and both transcription and genic orientation led us to hypothesize that only one of the two replicated gene copies is preferentially transcribed during S-phase.

We are now trying to better understand the molecular mechanisms responsible for chromatin maturation and to find out if and how chromatin maturation affects the transmission of transcription states to replicated gene copies. We have therefore combined ChIP with NChAP (ChIP-NChAP) to map the distribution of RNAPol2, histone H3, H3K56ac (new histones), and H3K4me^3^ and H3K36me^3^ (these marks of active transcription were used as proxies for old histones as in (*14*)) on replicated daughter genomes. In addition to wt yeast, our assay was carried out in mutants that have been linked to either new nucleosome assembly (rtt109Δ(*18*)) or old histone recycling (Mcm23A (*10, 15, 16*)). Using ChIP-NChAP, we were able to show that all examined histone marks and RNAPol2 are initially enriched on either the leading or the lagging strand copy depending on which strand replicated first. We found out that RNAPol2, H3K4me^3^ and H3K56ac enrichments shift to the other daughter chromatid as chromatin matures following replication. We also show that H3K56ac has no direct role in RNAPol2 binding to replicated gene copies. Likewise, the asymmetrical distributions of H3K4me^3^ and H3K36me^3^ correlate weakly or not at all with the asymmetrical distribution of RNAPol2 later in S-phase, suggesting that these histone marks do not carry or transmit epigenetic information on the transcriptional activity of the underlying gene and do not influence RNAPol2 (re)binding after replication. We have also uncovered hundreds of genes with an old nucleosome bias for the lagging gene copy in Mcm23A mutants, arguing against the recently advanced hypothesis that Mcm2 is directly involved in the recycling of old nucleosomes specifically to the lagging strand (*15, 16*).

We propose a two-step chromatin maturation model that explains how the relative differences in locus specific replication timing between the leading and the lagging daughter chromatid direct the asymmetrical distribution of new and old nucleosomes and RNAPol2 on replicated DNA. We show that chromatin and transcription states are inherited independently from each other. We propose that RNAPol2 is also recycled behind the fork. We suggest that this recycled RNAPol2 that shifts from the copy populated with old histones with “active” marks to its sister copy with new nucleosomes, is the true inherited epigenetic factor that puts “active” marks on new histones by transcribing that gene copy and recruiting methyltransferases. Active transcription states are therefore not restored by a copying mechanism that uses the chromatin state transmitted by the recycling of old nucleosomes as a template. We propose instead that the recycling of the transcription machinery itself which sequentially activates sister gene copies ensures the faithful transmission of active chromatin states and transcription states to both daughter chromatids after disruptions caused by the passage of the replication fork.

## Results

### RNAPol2 ChIP-NChAP

Replication doubles the gene copy number, which raises the question how the cellular transcription machinery transitions from regulating transcription on only one gene copy to dealing with the transcription of two copies? Does the transcriptional output increase two fold immediately after gene copy number doubling and are both gene copies transcribed equally? If so, RNAPol2 occupancy on replicated gene copies should also double shortly after or concomitantly with replication. On the other hand, if transcription outputs stay the same as before replication for a period of time as recently observed (*19*), a plausible explanation would be that the delay in the expected two-fold increase in transcriptional output is simply caused by limiting amounts of locally available RNAPol2. Gene expression can clearly not go up even if DNA content has doubled if local RNAPol2 levels are the same as before replication. We therefore first tested whether RNAPol2 is limiting after replication. We measured RNAPol2/DNA ratios for all S. Cerevisiae genes during S-phase using two channel DNA microarrays. The HA tagged Rpb3 subunit of RNAPol2, was immuno-precipitated from a synchronized cell population 25min (early S-phase) and 32min (early-mid S-phase) after release from α factor induced G1 arrest. The isolated DNA fragments from ChIP and input fractions were then amplified, labeled, mixed and hybridized to whole genome yeast DNA microarrays. We see that as more genes in the population are replicated, the ratios of RNAPol2 (ChIP DNA) to DNA content (i.e. input DNA) progressively decrease in replicated genes compared to non-replicated genes (**Figure S1A**). This confirms that RNAPol2 levels are indeed limiting after gene copy number doubling. In other words, the increase in locally available RNAPol2 amounts lags behind gene copy number doubling thus creating the recently observed “deficit” in mRNA production from replicated gene copies (*19*).

We next asked how those locally limiting RNAPol2 complexes divide between the two replicated gene copies. We used the RNAPol2 ChIP-NChAP assay to measure the differences in RNAPol2 enrichment between two newly replicated daughter chromatids. The strategy is to grow cells in the presence of the thymidine analogue EdU which labels newly replicated DNA strands and then perform ChIP of RNAPol2 with an HA tagged Rpb3 subunit. This is followed by the isolation of nascent DNA fragments from ChIP-ed DNA using streptavidin pull down after biotinylation of incorporated EdU. The purified nascent DNA is then used to make strand-specific deep-sequencing NChAP libraries as described previously (*17*). Thus, Watson and Crick strand sequencing reads obtained from these libraries originate from one or the other replicated daughter chromatid, respectively (**Figure 1A**). A comparison between the non-replicated ChIP (RNAPol2 on non-replicated chromatin, rows 8-9 from bottom), ChIP-NChAP (RNAPol2 on replicated chromatin, rows 10-11), and NChAP fractions (replicated chromatin, rows 12-13) from a synchronized population in early S-phase confirms that we are indeed able to isolate replicated DNA specifically bound by RNAPol2 (**Figure 1B**), as only RNAPol2 peaks from replicated regions (see the NChAP fraction) are enriched in the ChIP-NChAP fraction.

**Figure 1:**
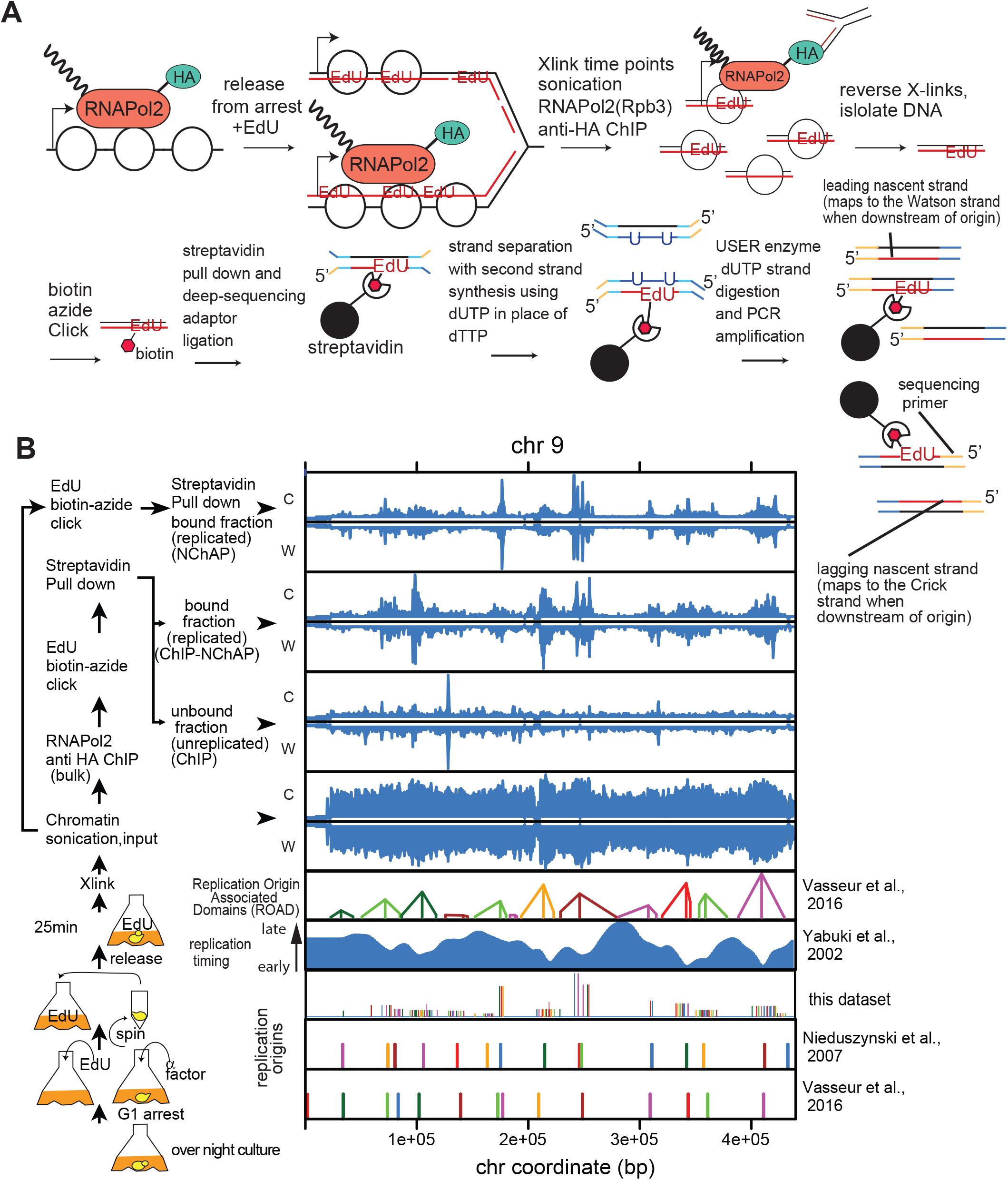
RNAPol2 ChIP-NChAP in early S-phase. **A.** Diagram of the RNAPol2 ChIP-NChAP experiment. **B.** RNAPol2 distribution on chromosome 9 from chromatin fractions diagramed on the left 25min (early S-phase) after release from G1 arrest (blue bars). The positions of replication origins (ARS- Autonomously Replicating Sequence) are shown in the three bottom rows: 1. previously documented ARS (Nieduszynski et al., 2007); 2. ARS identified in Vasseur et al, 2016, and 3. ARS from this study. NChAP from early S-phase in this study reveals clusters of replication origins at loci where only single origins were identified previously (rows 1, 2). The resolution of replication origin detection was increased in this dataset because all DNA fragments including the ones smaller than 100bp were kept for NChAP library construction unlike in Vasseur et al. (2016) which used mononucleosomal sized fragments (~150bp). Read counts from all fractions were grouped in 50 bp bins and first normalized to the genome average read count and then to the highest peak value in each chromosome. ROADs are Replication Origin Associated Domains i.e. regions that have been replicated from the same origin 25min after release as determined in Vasseur et al. (2016). W and C are Watson and Crick strand reads, respectively.

### Asymmetric Distribution of RNAPol2 on Daughter Chromatids

Is there a pattern in the distribution of RNAPol2 complexes on replicated gene copies? The median read density in the coding region of each gene (promoters excluded) from Watson (W) or Crick (C) reads was used as a measure of relative RNAPol2 occupancy on each gene copy (**Figure 2A**). The heat map in Figure 2A showing median read densities of all yeast genes from ChIP, NChAP and ChIP-NChAP fractions at different time points before and during S-phase (late G1 (2 replicates), early S (54% of the genome still not replicated), mid-early S (21% non-replicated), and early-mid S-phase (10% non-replicated, 2 replicates) confirms the specificity and reproducibility of our assay as RNAPol2 enrichment in the ChIP-NChAP fraction is only detected on replicated genes (compare ChIP-NChAP and NChAP fractions) and RNAPol2 occupancy correlates well with mRNA abundance (**Figure S1B**).

**Figure 2:**
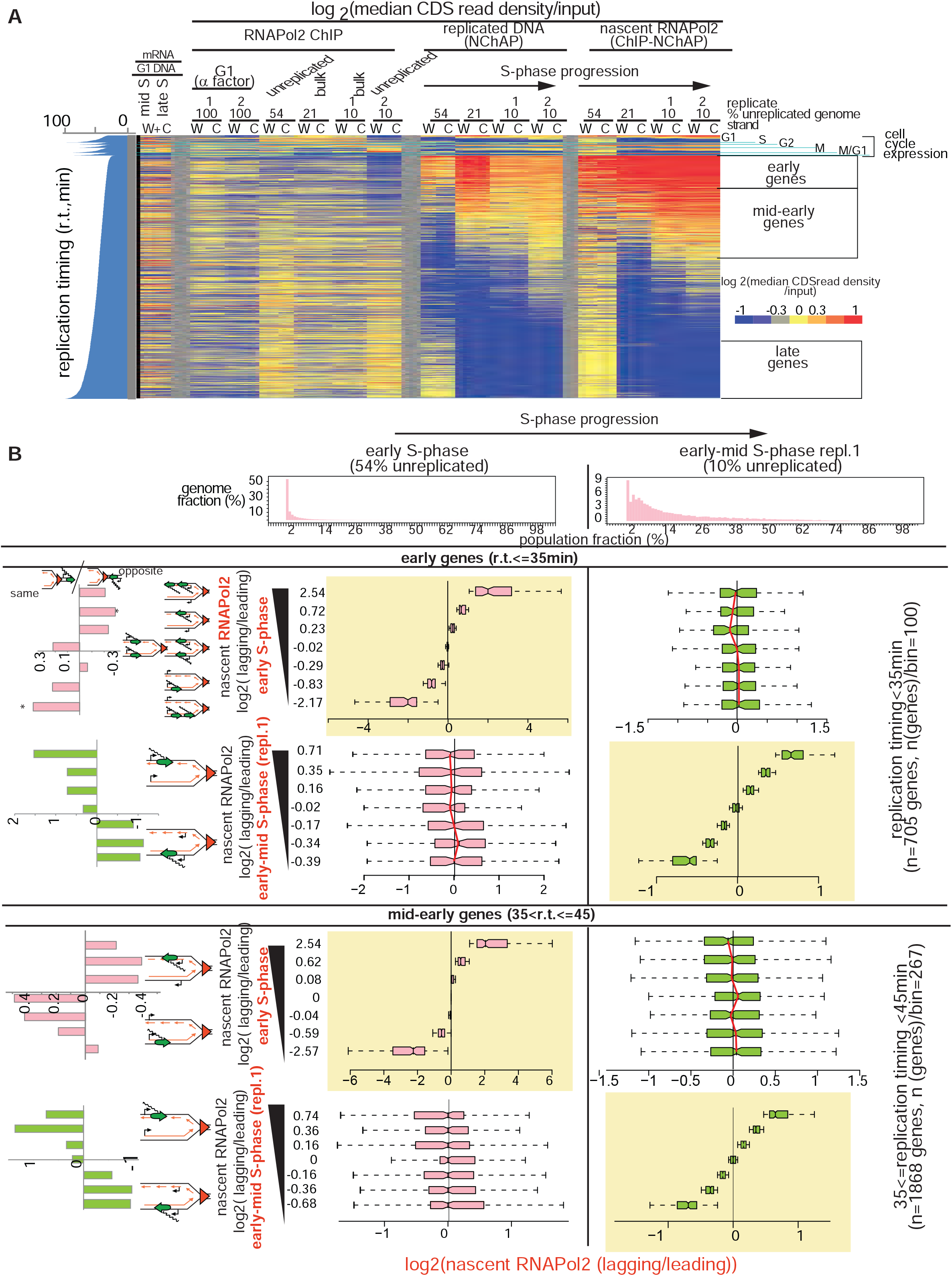
RNAPol2 is distributed asymmetrically on replicated gene copies. **A**. Heat map of median RNAPol2 occupancies in coding regions (CDS) of all yeast genes. Each line is an individual gene and columns represent occupancy values for (W)atson and (C)rick gene copies for different G1 and S-phase time points (late G1 (after 3.75 hrs in α factor, 100% of the genome is unreplicated, 2 replicates), early S (54% of the genome is unreplicated over the whole cell population), mid-early S (21% unreplicated), early-mid S (replicate 1, 10% unreplicated), and early-mid S (replicate 2, 10% unreplicated), from the ChIP, NChAP (nascent chromatin) and ChIP-NChAP (Nascent RNAPol2) fractions. The first two columns on the left represent mRNA enrichment over G1 genomic DNA in mid and late S (in the absence of EdU) determined with gene expression microarrays (Vasseur et al., 2016). Genes are grouped by cell cycle expression patterns and then ordered by replication timing (r.t.) within each group (r.t. from Vasseur et al., 2016). Median read density values for each gene have been normalized by separately dividing the W and C read densities of each gene in the ChIP and NChAP fractions with the W and C average read density of the sonicated input fraction for the same gene. The ChIP-NChAP medians have been normalized by dividing the W and C values with the input normalized W and C average values from the ChIP fraction. The unreplicated ChIP fractions from early (54% unreplicated) and mid early S (10% unreplicated, repl. 2) show RNAPol2 enrichment (yellow/red) in unreplicated genes (blue genes in the NChAP fractions) relative to replicated ones (blue/yellow), in contrast to bulk ChIP fractions from early-mid S (21% unreplicated), mid early S (replicate 1, 10% unreplicated). This indicates that we are successfully separating RNAPol2 bound nascent chromatin from RNAPol2 bound unreplicated chromatin **B.** Box plot distributions of lagging/leading nascent RNAPol2 ratios from early and early-mid S-phase (columns left to right, respectively) for early (1^st^ and 2^nd^ row from the top) and mid-early genes (rows 3 and 4) (r.t. = replication timing). The header row shows the distribution of genome read densities (in 400bp bins) normalized to the maximum read density for each NChAP fraction (reads have not been normalized to input) at indicated time points in S-phase. In early S-phase 54% of the genome has a read density of 0, i.e. 54% of the genome has not yet been replicated, and ~5% of the genome has a read density of 4, i.e. 4% of the population has replicated 5% of their genome. By early-mid S-phase 50% of cells have replicated at least 0.5% of their genomes and only 10% of the genome has not been replicated in the whole cell population. In the 1^st^ and 3^rd^ row from the top genes have been sorted by decreasing lagging/leading RNAPol2 occupancy in early S-phase and divided into 7 bins (y axis), and box plot distribution of nascent RNAPol2 lagging/leading ratios (x axis) have been determined for each bin in both time points. For example, early genes (705 genes shown in A) were sorted by increasing lagging/leading nascent RNAPol2 ratio from the early S-phase dataset and then divided into 7 bins of ~100 genes each, and we determined the box plot distribution of lagging/leading ratios for each bin. The average lagging/leading ratios for each bin are in the y axis on the left. For example the bottom group of genes in row 1 column 1 has on average 5.6 times more RNAPol2 on the leading copy than on the lagging. RNAPol2 distribution that was asymmetrical in early S-phase appears to even out later on in S-phase (rows 1 and 3, column 2). The bar graphs on the left show the enrichment of “same” genes for each bin indicated in the Y axis calculated as the ratio of “same” orientation genes versus “opposite” genes for each group normalized to the same/opposite ratio of all 705 early (rows 1 and 2) or 1868 mid-early genes (rows 3 and 4).

The scatter plot comparing median read densities of W and C gene copies of 705 early replicating genes shows that the early S-phase differences in RNAPol2 occupancy between two replicated gene copies are greater on nascent chromatin (the ChIP-NChAP fraction) than on non-replicated chromatin (ChIP fraction that did not bind to streptavidin beads). The observed asymmetry in RNAPol2 occupancy is also not due to a sequencing or EdU incorporation bias for one strand over the other because differences in W and C read densities are smaller in the nascent chromatin fraction (NChAP fraction) for which we use the same library construction protocol as for the ChIP-NChAP fraction (**Figure S1C**).

Next, we determined the pattern of RNAPol2 distribution between leading and lagging copies relative to genic orientation. Lagging and leading copy annotations for each W and C copy were assigned as described previously (*17*): W reads upstream of the closest replication origin (see Table S1 and Materials and Methods for replication origin mapping) originate from the lagging strand copy, while the complementary C reads are from the leading copy. The opposite is true for reads located downstream of origins. We calculated the ratio of RNAPol2 occupancies between the lagging and the leading gene copy for all 705 early genes in early S-phase, sorted the ratios from lowest to highest, and then divided the set into 7 bins of ~100 genes each (row 1 from top**, Figure 2B**).

The difference in RNAPol2 occupancy between lagging and leading gene copies appears to be greatest for genes with a low to moderate total RNAPol2 density (i.e. genes with low to moderate gene expression) (**Figure S1D**). This is to be expected if RNAPol2 and transcription factors (TFs) are recycled behind the replication fork and the binding of “new” RNAPol2 complexes and “new” TFs to replicated genes is limited at least in the early period after replication (**Figure S1A**). Small quantities of RNAPol2 that were bound to genes before replication are more likely to partition asymmetrically between the two replicated gene copies than large numbers of RNAPol2 complexes which are more likely to be distributed symmetrically.

The partitioning pattern of the locally available RNAPol2 complexes could then either be unbiased and stochastic or it could have an inherent bias i.e. a preference for the lagging or the leading copy. As shown in **Figure 2B**, RNAPol2 distribution has an apparent bias in ~50% of early replicating genes. Genes in the first bin (bottom bin, row 1, column 1 in **Figure 2B**) have on average 5.6 times more RNAPol2 on the leading copy and genes in the last bin (top bin) have 5.8 times more RNAPol2 on the lagging copy. These same gene bins that had a strong bias for one of the two copies appear to lose that bias later on in S-phase (compare columns 1 and 2 in rows 1 and 3 in **Figure 2B**). The RNAPol2 distribution bias in early-mid S-phase is now detected on different genes than in early S-phase (compare columns 1 and 2 in rows 2 and 4 in **Figure 2B**). We even record a slight anti-correlation between lagging/leading ratios in early S-phase and early-mid S-phase (row 4 in **Figure 2B**). In other words genes that have more RNAPol2 on the leading or the lagging copy in early-mid S-phase tended to have more RNAPol2 on the lagging or the leading copy in early S, respectively.

Next, we calculated the ratio of “same” orientation genes versus “opposite” genes for each bin in which genes were organized by increasing lagging/leading ratios of RNAPol2 occupancy in early S (column 1, rows 1 and 3, **Figure 2B**) or early-mid S (column 2, rows 2 and 4, **Figure 2B**) and normalized it to the “same”/”opposite” ratio of all 705 early (rows 1 and 2) or 1868 mid-early (rows 3 and 4) genes. As predicted from our nucleosome positioning maturation results (*17*), “same” gene enrichment is inversely proportional to the nascent RNAPol2 lagging/leading ratio in early S-phase, i.e. “same” genes in early S-phase tend to have more RNAPol2 on the leading copy and “opposite” genes tend to have more RNAPol2 on the lagging copy (rows 1 and 3, **Figure 2B**). Later on in early-mid S-phase, the observed RNAPol2 “polarity” appears to switch and RNAPol2 becomes more abundant either on lagging copies when transcription and replication travel in the same direction or on leading copies when they are opposite (rows 2 and 4**, Figure 2B**). Taken together, the switch in genic orientation “preference” and the apparent anti-correlation of RNAPol2 occupancy bias between early and early-mid S-phase suggest that RNAPol2 binds to one replicated gene copy first and then switches to the other later.

This RNAPol2 enrichment bias is specific to nascent chromatin as we do not observe significant differences in RNAPol2 occupancies on lagging and leading strands in bulk or non-replicated chromatin (**Figure S2**). More importantly, the asymmetrical pattern of RNAPol2 distribution observed on early and mid-early replicating genes in early-mid S-phase is highly reproducible (**Figures S3 and S4**). We detect the same RNAPol2 distribution pattern within the same gene bins in four different biological replicates (2 are shown in Figures 2A and S3, and the other 2 in Figure S4B, row 4), suggesting that this “later” RNAPol2 configuration represents the final step in the process of RNAPol2 binding to replicated genes.

The RNAPol2 distribution pattern in early-S phase is on the other hand difficult to reproduce since most genes are on the first very transient step of the RNAPol2 (re)binding process. Due to the stochastic nature of replication origin activation and non-synchronous fork progression in different cells in the population, the same genes from different early-S phase replicates (even though they appear to be at comparable points in the replication program) are at slightly different points relative to the position of their respective replication fork. Consequently RNAPol2 lagging/leading gene ratios do not correlate in early S-phase replicates (compare columns 1 and 2 in **Figure S4B**). We observe nevertheless similar trends in the RNAPol2 distribution pattern in the two replicates: prevalence of RNAPol2 on the leading copy of “same” genes and on the lagging copy of “opposite” genes in early S-phase.

### The asymmetric distribution of RNAPol2 on daughter chromatids is independent of the asymmetric distribution of new histones

Next we asked if the pattern of RNAPol2 binding to replicated gene copies correlates with and is dependent on the pattern of nucleosome binding. We measured the dynamics of new nucleosome binding first. Acetylation of Lysine 56 on Histone H3 with the Rtt109 histone acetyl-transferase, marks newly synthesized histones in yeast (*18, 20*). It is consequently enriched at promoters with high H3 turnover rates and on newly replicated DNA (*18*). So, we checked whether H3K56ac distribution on daughter chromatids correlates with the asymmetric distribution of RNAPol2 described above.

H3K56ac ChIP-NChAP from a synchronized cell population in mid-S-phase shows that the distribution of “new” histones indeed correlates with the RNAPol2 distribution from early-mid S-phase (**Figure 3**, compare columns 3 an 4 in rows 3 and 4 from top): RNAPol2 is enriched on the gene copy that also contains more new acetylated histones. H3K56ac and RNAPol2 lagging/leading ratios do not correlate in early S-phase and even appear to be somewhat anti-correlated (columns 1 and 2, rows 3 and 4 from top in **Figure 3**). The lack of correlation between the distribution patterns of new histones and RNAPol2 in early S-phase was also confirmed in a replicate experiments where H3K56ac and RNAPol2 ChIP-NChAPs were performed in parallel from the same cell culture in each replicate (left and right panels, column 2, rows 2 and 3 from top, **Figure S3B**). H3K56ac does however have a similar bias for “same” leading and “opposite” lagging gene copies as RNAPol2 in early S-phase, albeit on different genes (compare rows 2 and 3 from the top in **Figure S3 B**).

**Figure 3:**
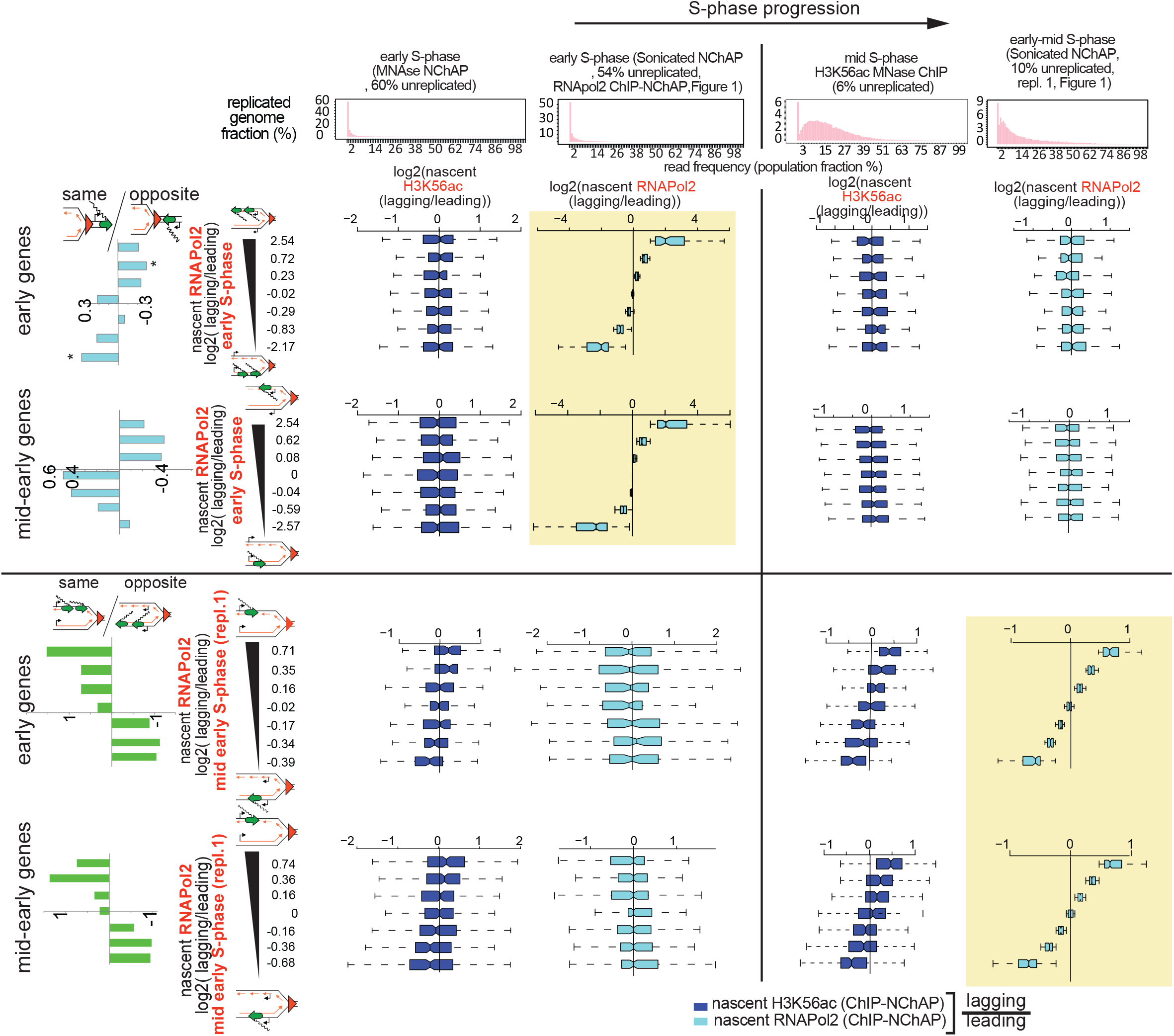
The asymmetric distribution of RNAPol2 on daughter chromatids is independent of the asymmetric distribution of new histones. Box plot distributions of lagging/leading nascent H3K56ac (dark blue) and nascent RNAPol2 (light blue) ratios from early (columns 1 and 2, respectively) and mid and early-mid (columns 3 and 4, respectively) S-phase for early (1st and 3^rd^ row from top) and mid-early genes (2^nd^ and 4^th^ row from top). Header: The distribution of genome read densities (in 400bp bins) normalized to the maximum read density for each dataset as a measure of S-phase progression as in Figure 2B. From left: 1. nascent chromatin from early S-phase (MNase-NChAP, 60% non-replicated); 2. nascent chromatin early S-phase replicate 1 from Figure 2 (54% non-replicated); 3. bulk H3K56ac ChIP from mid S-phase (6% non-replicated, since H3K56ac is a mark of new histones that area incorporated into replicated DNA the read density of the H3K56ac ChIP mid-S fraction mirrors DNA replication and can serve as a proxy for measuring what fraction of the genome has replicated in the cell population); 3. nascent chromatin mid-early S-phase replicate 1 from Figure 2 (10% non-replicated). Reads have not been normalized to input. By mid S-phase 15% of cells have replicated at least 3% of their genomes and only 6% of the genome has not been replicated in the whole cell population. Rows 1 and 2: early and mid-early genes have been sorted by increasing lagging/leading RNAPol2 occupancy from early S-phase (Figure 2), respectively, and then divided into 7 bins as in Figure 2B (y axis), and box plot distributions of nascent H3K56ac lagging/leading ratios (x axis) from early (left) and mid (right) S-phase have been determined (dark blue boxes) and compared to nascent RNAPol2 lagging/leading ratios from early and early-mid S-phase (light blue boxes). The bar graphs on the left show the “same” gene enrichment calculated as in Figure 2B for gene bins indicated in the Y axis of each row on the right. Rows 3 and 4: as rows 1 and 2 but sorted by increasing lagging/leading RNAPol2 occupancy from early-mid S-phase (replicate 1, Figure 2).

The lack of correlation between RNAPol2 and H3K56ac lagging versus leading enrichment ratios in early S-phase associated with a similar dependence on genic orientation for the distribution pattern of H3K56ac and RNAPol2 suggests that new nucleosomes and RNAPol2 follow the same order of binding to daughter chromatids but that the two processes are independent of each other. The coincidence of H3K56ac and RNAPol2 enrichments later in S-phase is consequently due to the convergence of these two independent pathways. We therefore conclude that H3K56ac does not influence or direct RNAPol2 binding on newly replicated genes. Indeed, RNAPol2 follows the same asymmetric distribution pattern that correlates with genic orientation and that appears to switch from one gene copy to the other even in the absence of H3K56ac in rtt109Δ cells (**Figure 4B**). A duplicate RNA-seq experiment using spike-in normalization with total RNA from *S.pombe*, shows a ~30% genome wide reduction of mRNA levels in rtt109Δ mutants compared to wt cells. It is therefore unlikely that H3K56ac directly suppresses transcription of newly replicated genes as proposed recently (*19*) (**Figure S5**).

**Figure 4:**
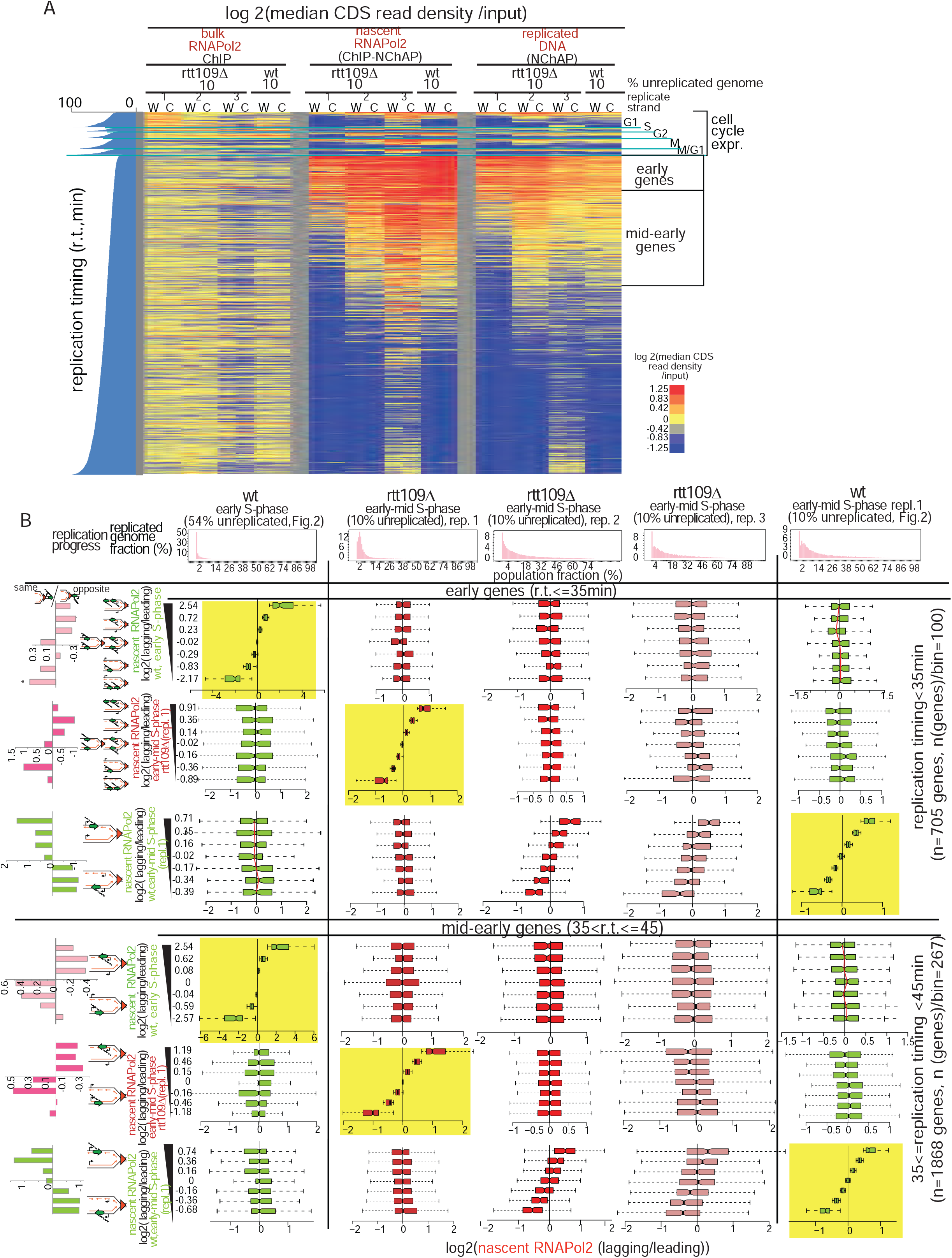
RNAPol2 is distributed asymmetrically in the absence of H3K56ac in rtt109Δ cells. **A**. Heat map of median RNAPol2 occupancies in coding regions (CDS) of all yeast genes. Each line is an individual gene and columns represent occupancy values for (W)atson and (C)rick gene copies for early-mid S-phase after release from G1 arrest in rtt109Δ(10% non-replicated,3 biological replicates) and wt (10% non-replicated replicate 1 from Figure 2), from the ChIP, NChAP (nascent chromatin) and ChIP-NChAP (Nascent RNAPol2) fractions. Genes are grouped as in Figure 2A. Median read density values for each gene have been normalized as in Figure 2A. **B.** Box plot distributions of lagging/leading nascent RNAPol2 ratios from early (wt) to early-mid S-phase (wt and rtt109Δ, columns left to right) for early (1st to 3rd row from the top) and mid-early genes (three bottom rows) (r.t.= replication timing). Header: distribution of genome read densities (in 400bp bins) normalized to the maximum read density for each NChAP fraction (reads have not been normalized to input) at indicated time points in S-phase as in Figure 2. Rows 1 and 4 from the top: genes have been sorted by increasing lagging/leading RNAPol2 occupancy in early S-phase (wt, Figure 2B) and divided into 7 bins as in Fig.2 B (y axis), and box plot distributions of nascent RNAPol2 lagging/leading ratios (x axis) have been determined for each bin at indicated time points. Rows 2,5 and 3,6: same as 1 and 4 except that genes have been ordered by increasing lagging/leading RNAPol2 ratios from rtt109Δ (replicate 1) or early-mid S (replicate 1, wt Fig. 2), respectively. The bar graphs on the left show “same” gene enrichments calculated as in Figure 2B for gene bins indicated in the Y axis of each row on the right. The RNAPol2 distribution pattern between leading and lagging strand gene copies in rtt109Δ cells from replicates 2 and 3 correlates with wt (early-mid S, replicate 1 from Figure 2) indicating that the asymmetric distribution of RNAPol2 on replicated DNA is independent of the distribution of H3K56Ac.

### “Old” and “New” Histone and RNAPol2 distributions on replicated gene copies are determined by local differences in the timing of leading and lagging strand replication

Our next question was whether the distribution pattern of old histones influences RNAPol2 binding to replicated genes. In order to establish a comprehensive timeline of nucleosome assembly on replicated DNA and explore how the reassembly of old nucleosomes influence RNAPol2 distribution on replicated DNA we performed parallel ChIP-NChAP experiments for H3, H3K4me^3^ and H3K36me^3^ (these methylated histones were used as “proxies” for old histones as in (*14, 15*)), H3K56ac and RNAPol2 in two biological replicates at 20 and 25 min after release from G1 arrest (**Figures 5, S6A-C** and **S7A**).

**Figure 5:**
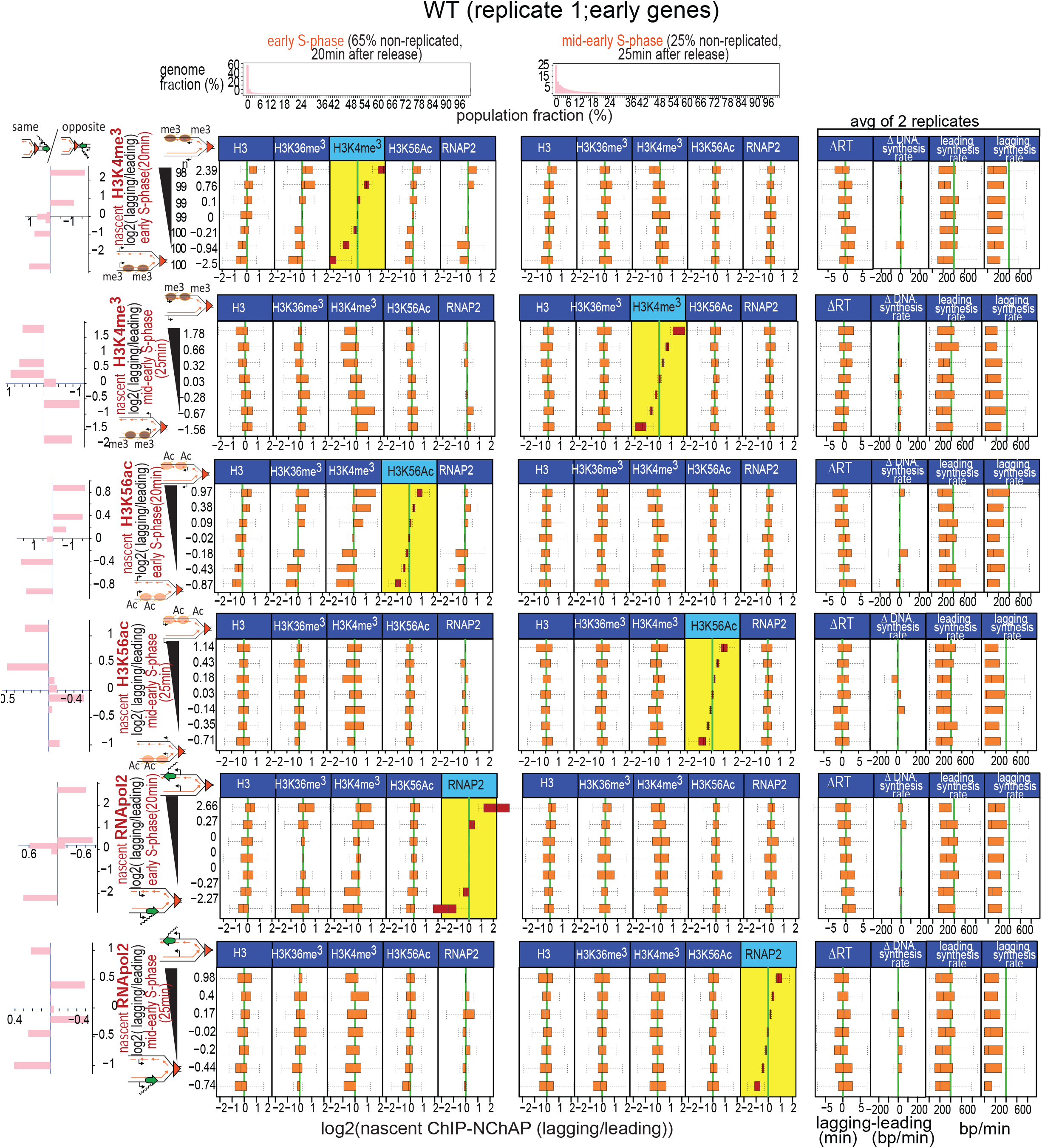
Old nucleosomes are recycled to the daughter chromatid that replicated first. Box plot distributions of lagging/leading ChIP-NChAP ratios for H3, H3K36me3, H3K4me3, H3K56ac and RNAP2 from early and mid-early S-phase for early genes measured in the same culture of wt cells (biological replicate 1). The histograms in the header show the distribution of genome read densities (in 400bp bins) normalized to the maximum read density for each NChAP (replicated DNA) fraction (reads have not been normalized to input) at indicated time points in S-phase. In early S-phase (20min time point) 65% of the genome has a read density of 0, i.e. 65% of the genome has not yet been replicated. By mid-early S-phase (25min time point) 25% of the genome has not been replicated in the whole cell population. Rows 1-6: genes have been sorted by decreasing lagging/leading occupancy of H3K4me3 in early S, H3K4me3 in mid-early S, H3K56ac in early S, H3K56ac in mid-early S, RNAP2 in early S and RNAP2 in mid-early S, respectively (yellow background), and divided into 7 bins (y axis on the left). Box plot distribution of lagging/leading ratios (x axis) for the chromatin features indicated in the headers have been determined for each bin. For example, in row 1 early genes have been sorted by decreasing lagging/leading early S H3K4me3 ratios and then divided into 7 bins of ~100 genes each (n on the left in row 1), and we determined the box plot distribution of lagging/leading ratios for H3, H3K36me3, H3K4me3, H3K56ac and RNAP2 in early and mid-early S for each bin (average lagging/leading early S H3K4me3 ratios for each bin are shown on the y axis on the left). Thus, the bottom group of genes has on average 5.6 times more H3K4me3 on the leading copy than on the lagging. We then calculated the ratio of “same” orientation genes versus “opposite” genes for gene bins in each row normalized to the same/opposite ratio of all 705 genes (bar graphs, column 1, left). The bar graphs show the “same” gene enrichment for gene bins indicated in the Y axis of each row on the right. Columns 11-16: box plot distributions of 11- ΔRT: the difference in replication timing (RT) between the lagging and the leading strand for each gene (RT for each gene is the median RT of all 50bp segments in the CDS of the gene determined from two replicate time-courses in Figure S8); 12- ΔDNA synthesis rate: average difference between lagging and leading DNA synthesis rates for each gene in the bin. Synthesis rates were calculated as in Figure S8 using replication timing from Figure S8 and ROADs determined from NChAP fractions of the 20 et 25min time points of two biological replicates (replicate 1 from Figure 5 and replicate 2 from Figure S7); 13 and 14- average leading and lagging DNA synthesis rates used to obtain the Δ DNA synthesis rate in column 12.

Our experiments show that H3K4me^3^ and H3K36me^3^ distributions follow a similar pattern to the one seen for H3K56ac (**Figures 5, S6C and S7A**). In early S-phase (20min after release from arrest), H3K4me^3^ and H3K36me^3^ have a bias for either the leading or the lagging copy in a comparable number of genes and this bias correlates with the binding patterns of H3, H3K56ac and RNAPol2 (**Figure 5**).

Also, as recorded above for H3K56ac and RNAPol2, “same” gene enrichment is inversely proportional to the nascent H3/H3K36me^3^/H3K4me^3^/H3K56ac/RNAPol2 lagging/leading ratio in early S-phase in both biological replicates (**Figures 5 and S7A**). In other words, “same” genes tend to have more H3/H3K36me^3^/H3K4me^3^/H3K56ac/RNAPol2 on the leading copy and “opposite” genes tend to have more of these chromatin features on the lagging copy in early S-phase (**Figure 5**). As seen earlier for RNAPol2 and H3K56ac, the genic orientation bias switches for H3K4me^3^ as well: “same” genes have more H3K4me^3^/H3K56ac on the lagging strand and “opposite” genes have more H3K4me^3^/ H3K56ac on the leading strand in mid-early S (25min time point) of the first biological replicate. As chromatin matures after replication, we observe the same switch in occupancy for H3K4me^3^ as for H3K56ac and RNAPol2 above. H3K4me^3^ shifts either from the leading to the lagging gene copy at “same” genes or from the lagging to the leading copy at “opposite” genes. This switch can be directly observed for H3K4me^3^ in **Figure 5** where the H3K4me^3^ lagging/leading ratios from mid-early S-phase are anti-correlated to the H3K4me^3^ lagging/leading ratios from early-S-phase. Remarkably, the 5’ enrichment of H3K4me^3^ and 3’ enrichment of H3K36me^3^ that is characteristic of coding regions of actively transcribed genes is already apparent in the earliest time point. This is consistent with an earlier finding that old histones are recycled close to their original locus before replication (*21*) (**Figure S6 C**). Note that this polarity shift was not observed in the second biological replicate. This is because the 25min time point in the second replicate represents an earlier stage of S-phase when 39% of the genome is still not replicated compared to the replicate in Figure 5 where only 25% of the genome has not yet replicated (**Figure S7A**). Consequently, the 20 and 25min points of the second replicate have essentially the same H3/ H3K36me^3^/ H3K4me^3^/ H3K56ac/ RNAPol2 distribution pattern between lagging and leading copies.

At 25min after release from arrest, chromatin maturation has progressed further in the first replicate than in the second replicate. Thus at this more advanced stage of chromatin maturation in the first replicate, the correlation in occupancy bias between all measured chromatin features that was observed in early-S phase “disappears” in the later time-point. All chromatin features in mid-early S-phase appear to be symmetrical except for the feature that was used to sort genes into bins of increasing lagging/leading ratios. In other words, we do detect equal numbers of genes with significantly asymmetrical partitioning of all measured chromatin features including H3 and H3K36me3 (not shown) and not just RNAPol2, H3K4me3 and H3K56ac that are shown in Figure 5 but the asymmetrical distribution of any one of these features is not found on the same genes as the asymmetrical distribution of any other feature.

The asymmetrical partitioning of histones and RNAPol2 on newly replicated daughter chromatids can be detected on about half of early (**Figure 5**) and mid-early replicating genes. These asymmetrical distribution patterns cease to correlate as chromatin matures. This asymmetry is not equally “visible” for all chromatin features on the same genes because replication fork progression and chromatin maturation are not synchronized at the cell population level. The onset of replication from any origin is a stochastic process that happens within a limited probabilistic time window but nevertheless occurs at different times in different cells. Likewise, local fork velocities and chromatin maturation kinetics vary from cell to cell. Since the occupancy shift from one daughter chromatid to the other occurs independently for histones and RNAPol2, by mid-early S-phase different chromatin features will have matured at different times on the same genes in different cells. In some cells histones will be on the first step of nucleosome binding to replicated DNA and preferentially occupy the leading or the lagging strand, depending partially on genic orientation. Meanwhile, in other cells, the histone occupancy shift to the sister strand will have already taken place. The entire cell population will consequently consist of a mixture of cells with lagging and leading gene copies that have more old or new histones than their sister copy. The distribution of log_2_(lagging/leading) ratios for old or new histones in such a population will then have a median of 0 as we have observed. Even though RNAPol2 and histones follow the same steps when they bind to replicated gene copies, they are doing so independently of each other and with different kinetics. We consequently end up measuring apparently symmetrical histone distributions on genes that have asymmetrical RNAPol2 distributions and vice versa, symmetrical RNAPol2 distributions on genes that have asymmetrical histone distributions.

It is important to emphasize at this point that our results clearly show that the initial correlation between RNAPol2 and H3K4me^3^/H3K36me^3^/H3K56ac/H3 binding patterns in early S-phase is coincidental because that correlation disappears later in S-phase. In other words, the pattern of H3K4me^3^ and H3K36me^3^ recycling on replicated DNA does not dictate where RNAPol2 will bind. This is direct evidence that H3K4me^3^ and H3K36me^3^-long thought of as “epigenetic” marks of active transcription, merely because they are enriched on actively transcribed genes- are actually not epigenetic marks in the strict sense. Their “inheritance” to replicated gene copies does not have an effect on RNAPol2 occupancy and thus also has no role in the post-replicative re-establishment of transcriptional activity of their underlying genes. Consequently, H3K4me^3^ and H3K36me^3^ do not transmit information on the transcriptional state of genes prior to replication to their replicated gene copies.

We have by now seen that old histones can either be recycled asymmetrically with a preference for the lagging or the leading gene copy or symmetrically on both gene copies. Note that it is impossible to distinguish between symmetrical histone distribution in the same cell and asymmetrical distribution in the cell population with equal numbers of cells with a lagging or leading bias, due to the nature of ChIP-NChAP experiments which, as any next generation sequencing experiment using a population of cells, measure average occupancy in the population.

What determines how old histones will be distributed on replicated gene copies? The “choice” of the leading or the lagging gene copy in gene bins with a significant bias correlates strongly with genic orientation, i.e. with the direction of transcription relative to the direction of the replication fork. We therefore conjectured that the recycling of old histones will be influenced by local replication dynamics of the leading and lagging strand, which are in turn influenced by the interactions of the replication fork with the transcription machinery. We therefore measured leading and lagging replication timing of all genes as described in **Figure S8 A-E**.

Replication timing of genes that are replicated by the same replication fork i.e. on the same ROAD (Replication Origin Associated Domain, determined as described in Materials and Methods), gradually increases as genes get further away from the origin. We can consequently use these differences in replication timing between genes on the same ROAD to calculate average lagging and leading DNA synthesis rates on each replicated gene. Since DNA synthesis rates on any given gene are directly dependent on the average fork velocity at that locus we can use DNA synthesis rates as a measure of average fork speed through that gene. If we plot replication timing values separately for the leading and lagging strands versus the chromosomal coordinates of genes within the same ROAD upstream and downstream of each origin, we can estimate replication fork velocity from the slope of the linear fit for every replicated gene copy (**Figures S8 F-H and S6B**). It is then straightforward to calculate the differences in replication timing and DNA synthesis rates between lagging and leading gene copies of all replicated genes (**Figures S8 F-H and S6B**).

Our replication timing measurements reveal a remarkable trend that explains the observed pattern of nucleosome binding to newly replicated DNA (**Figure 5**). Old and new nucleosomes and RNAPol2 simply bind first to the gene copy that replicated earlier than its sister. If the leading gene copy has an earlier (lower) replication time than the lagging copy of the same gene, old and new nucleosomes and RNAPol2 will bind to that copy first. Conversely, if the lagging copy replicates before the leading copy, the lagging copy will be “chromatinized” first. The bias in total H3 distribution for the strand that replicated first supports our hypothesis that the strand that replicated later is “under-chromatinized” in this early chromatin maturation step. If the difference in replication timing is insignificant nucleosomes and RNAPol2 will randomly go to one or the other or to both copies resulting in an average log_2_(lagging/leading) ratio of 0. Surprisingly, even though lagging DNA synthesis rates are on average slower than the leading DNA synthesis rates in the gene population as a whole, as one might expect, the differences between leading and lagging synthesis rates on any individual gene are small and do not appear to influence where chromatin will be assembled first. We do observe however that leading strand DNA synthesis rates are overall faster than lagging rates on genes that favor leading strand nucleosome assembly in early S, while lagging strand synthesis rates are somewhat faster than leading rates on genes that are biased for the lagging strand (**Figures 5 and S7A**). Genic orientation is however, still a better predictor of which copy is more likely to replicate first and thus direct the order of chromatin assembly and RNAPol2 binding after replication. “Opposite” orientation genes tend to favor lagging replication over leading, while the leading gene copy replicates before the lagging copy for genes where transcription travels in the “same” direction as the replication fork.

We propose the following two step model for chromatin maturation. In step one, old nucleosomes (or at least their H3-H4 tetramers) with H3K4me^3^ and H3K36me^3^ are recycled very close behind the fork, which explains why the typical pattern of H3K4me^3^ and H3K36me^3^ distribution on coding regions is largely preserved on newly replicated genes. Old nucleosomes preferentially bind to the daughter chromatid that is replicated first: leading for “same” genes or lagging for “opposite” genes. If both chromatids are replicated almost simultaneously old nucleosomes and new nucleosomes will bind to one or the other chromatid randomly and in equal proportions. When the difference in replication timing between the two chromatids is significant, new histones and old histones will compete for binding to the only chromatid that has completed its synthesis and since the local concentration of old nucleosomes is higher than the local concentration of new nucleosomes, that daughter chromatid will end up with a nucleosome population consisting mostly of recycled old nucleosomes. RNAPol2 will at this point probably also be recycled from ahead of the fork and will also end up on the chromatid that replicated first or randomly on one or the other if they replicated nearly simultaneously. New nucleosomes will bind to the sister chromatid that replicated later in step 2 because most old nucleosomes have already been recycled to the chromatid that replicated earlier in step 1. RNAPol2 will then shift to the second daughter chromatid and direct Set1 and Set2 to respectively methylate new H3 histones on K4 and K36.

### The synchronicity of globally slower replication forks in Mcm23A mutants brings into focus the genome wide asymmetry of old histone recycling

Since the binding preferences of RNAPol2 and nucleosomes for the leading or lagging strands depend on their respective order of replication, we wanted to further explore the mechanism that regulates the replication timing of each strand. We decided to measure RNAPol2 and nucleosome binding to replicated DNA in Mcm23A mutant cells. We surmised that a mutant of Mcm2 - a subunit of the replication fork helicase Mcm2-7- was likely to affect replication fork progression. Moreover, recent studies have hypothesized that Mcm2 acts as a histone chaperone that is responsible for the transfer of old histones specifically to the lagging strand. This hypothesis was based on observations that alanine substitutions of tyrosines 79, 82 and 91 in the Mcm23A mutant impair the interaction of Mcm2 with histones ahead of the replication fork (*10*) and cause an apparent increase in asymmetric recycling of old histones with a bias for the leading daughter chromatid (*15, 16*).

Our results reveal however that the general trends for the distribution of new and old histones and RNAPol2 are remarkably similar in wt and Mcm23A cells. The difference lies mostly in the magnitude of asymmetrical nucleosome recycling. While almost all genes in the cell population appear to have more or less highly asymmetric histone distributions in the Mcm23A mutant “only” ~50% of gene copy pairs in wt cells are “asymmetrical”. The number of genes with a leading strand bias is nevertheless comparable to the number of genes with a lagging strand bias in the mutant and wt alike. This is not consistent with the hypothesis that Mcm2 recycles old histones specifically to the lagging strand as suggested recently (*15, 16*) (**Figures 6, S6D and S7B**). We observe instead that the dynamics of histone and RNAPol2 distribution on replicated daughter chromatids follow a similar pattern that has different kinetics in Mcm23A and wt cells. The positive correlation in the distribution of H3K4me^3^, H3K36me^3^ and H3 persists for a longer period in the mutants compared to wt and is still apparent even when 80% of the genome has been replicated in the whole population (**Figure 6A**). Unlike in wt cells, H3K56ac is already anti-correlated with the H3 methyl marks from the earliest time point in the Mcm23A mutant. We also do not observe the shift in H3K4me^3^ enrichment from one strand to the other that we detected in wt cells (replicate 1 from **Figure 5**).

**Figure 6:**
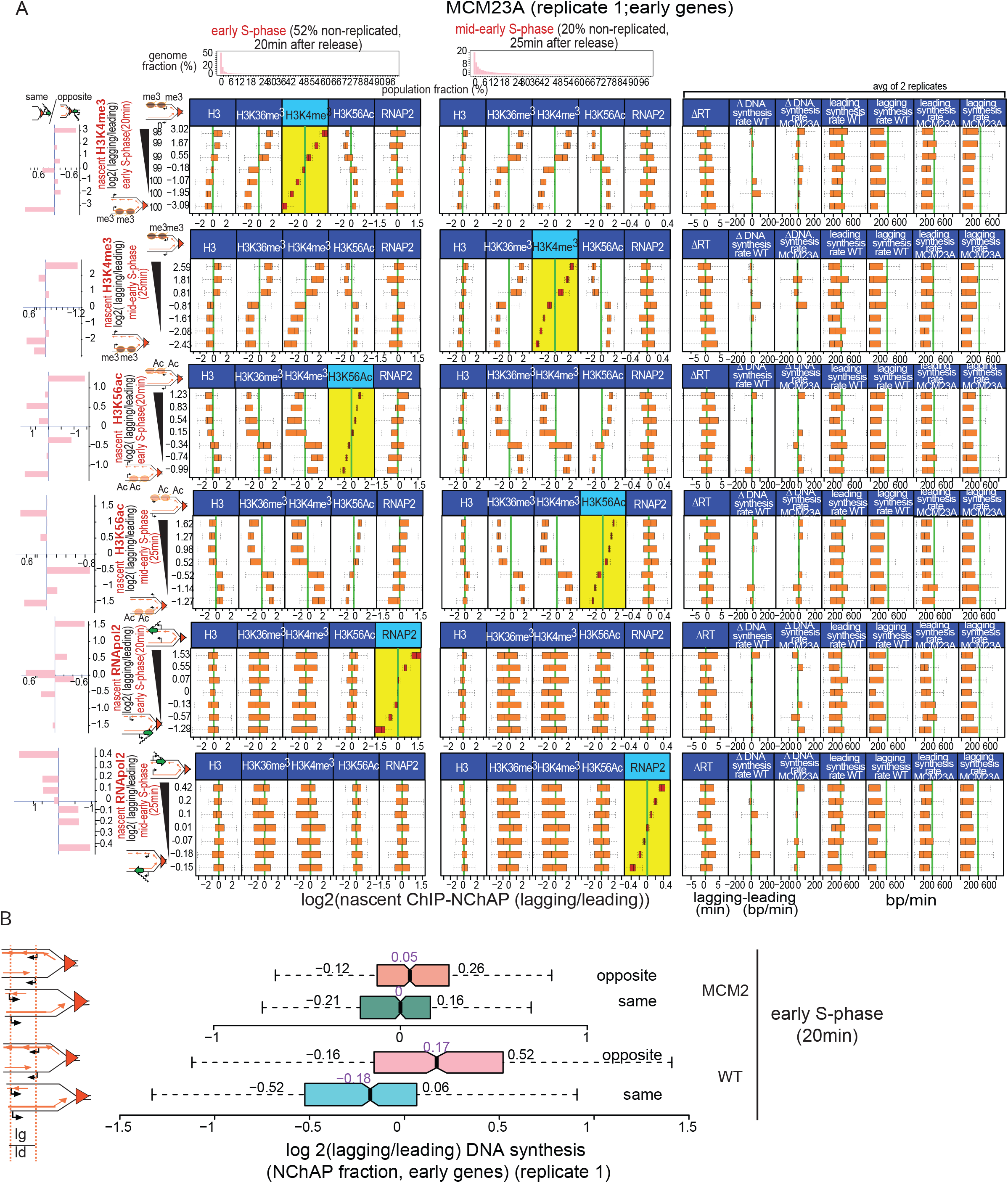
Chromatin maturation in Mcm23A cells. A. Box plot distributions of lagging/leading ChIP-NChAP ratios for H3, H3K36me3, H3K4me3, H3K56ac and RNAP2 from early and mid-early S-phase for early genes measured in the same culture of Mc23A mutant cells (biological replicate 1). The histograms on top show the distribution of genome read densities (in 400bp bins) normalized to the maximum read density for each NChAP (replicated DNA) fraction (reads have not been normalized to input) at indicated time points in S-phase. In early S-phase (20min time point) 52% of the genome has a read density of 0, i.e. 52% of the genome has not yet been replicated. By mid-early S-phase (25min time point) 20% of the genome has not been replicated in the whole cell population. Rows 1-6: genes have been sorted by decreasing lagging/leading occupancy of H3K4me3 in early S, H3K4me3 in mid-early S, H3K56ac in early S, H3K56ac in mid-early S, RNAP2 in early S and RNAP2 in mid-early S, respectively (yellow background), and divided into 7 bins (y axis on the left). Box plot distribution of lagging/leading ratios (x axis) for the chromatin features indicated in the header have been determined for each bin as in Figure 5. The bar graphs on the left show the “same” gene enrichment for gene bins indicated in the Y axis of each row on the right. Columns 11-19: box plot distributions of 11- ΔRT: the difference in replication timing (RT) between the lagging and the leading strand for each gene in Mcm23A cells (RT for each gene is the median RT of all 50bp segments in the CDS of the gene determined from the two replicate time-courses in Figure S8); 12- ΔDNA synthesis rate in WT: average difference from two replicates and two time points between lagging and leading DNA synthesis rates for each gene in the bin. Synthesis rates were calculated as in Figure S8 using replication timing from Figure S8 and ROADs determined from NChAP fractions of the 20 et 25min time points of two biological replicates (replicate 1 from Figure 6 and replicate 2 from Figure S8); 13- ΔDNA synthesis rate in Mcm23A; 14 and 15- average leading and lagging DNA synthesis rates in WT cells used to obtain the ΔDNA synthesis rate in column 12; 16 and 17- average leading and lagging DNA synthesis rates in Mcm23A cells used to obtain the ΔDNA synthesis rate in column 13. **B.** box plot distribution of DNA synthesis bias (log2(lagging/leading) of the NChAP fraction) for early replicating “opposite” and “same” genes in early S-phase (20min after release from G1 arrest) for wt replicate 1 (Figure 5) and Mcm23A replicate 1 (Figure 6A).

RNAPol2 occupancy on the other hand, does shift from one strand to the other between early and mid-early S-phase. The correlation with histone binding patterns is weak and heterogeneous: on some genes the RNAPol2 binding bias correlates better with H3K4me^3^/H3K36me^3^/H3 and on others it is better correlated with H3K56ac (compare rows 1 and 2 from the top to rows 3 and 4 in early S-phase in **Figure 6A**). RNAPol2 binding on genes with the largest bias for one or the other daughter chromatid does not correlate at all with the binding pattern of old or new histones in replicate 1 (**rows 5 and 6 from the top, Figure 6**) and has a weak correlation with H3K56ac and a weak anti-correlation with H3K4me^3^ and H3K36me^3^ in replicate 2 (**rows 5 and 6 from top in Figure S7B**). This is further evidence that RNAPol2 binding to newly replicated DNA is independent of H3K4me^3^, H3K36me^3^ or H3K56ac occupancy and that any correlation or anti-correlation between the RNAPol2 and histone binding patterns is coincidental and not causative.

As with wt cells, the determining factor in the “choice” of daughter chromatid to which old nucleosomes are preferentially recycled in Mcm23A mutants is replication timing. Consistent with the timeline we described above for wt cells: old nucleosomes bind first to the strand that replicated earlier and new nucleosomes then bind to the sister strand that replicated later. Genes in the mutant whose transcription goes in the same direction as the replication fork tend to replicate the leading strand first and “opposite” genes tend to replicate the lagging strand first, as in wt cells (**Figure S9D**). This is corroborated by the lagging/leading strand ratios of replicated DNA (NChAP fraction) for “same” and “opposite” genes in early S-phase (**Figure 6B**): lagging gene copies are more replicated than their leading sisters on opposite genes and leading copies are on average more replicated than their lagging counterparts on same genes. The trend is the same in wt and mutant cells.

Why does the mutant have more genes with an asymmetrical nucleosome distribution than the wt? Our replication timing measurements reveal that mid-late and late genes in the mutant replicate later than in wt cells (**Figure S8 D-E**). This is probably linked to the gradual slowing down of DNA synthesis rates in the mutant (**Figures S6B and S8H**). Indeed DNA synthesis rates in wt cells are similar for early genes and late genes (~340bp/min on average), while they are markedly slower for late genes relative to early genes in the mutant (~270bp/min for early genes compared to ~200bp/min for late genes). Local DNA synthesis rates, which are directly related to local replication fork velocities, are between 200 and 800 bp/min for most genes in wt and between 100 and 500bp/min for most genes in mutant cells. Our fork velocity estimates in wt cells are lower than the previously estimated velocity of 1 to 2kb/min (*22, 23*). Our calculations are based on synthesis rates within gene bodies while the previously published estimates are based on genome-wide replication rates. This may explain the discrepancy as replication forks are likely to be slowed down by the transcription machinery when they advance through genes. Since fork progression rates are globally slower in the Mcm23A mutant, we propose that the apparent persistency of asymmetrical histone distribution in the Mcm23A till late in S-phase is at least partially a reflection of better synchronization of Mcm23A replication forks in different cells of the population. In other words, because replication forks are globally slower in the mutant, they are more likely to be close to the same chromosomal location in different cells even though the forks in different cells started from origins that “fire” stochastically at slightly different times just as in wt cells. Consequently, replication forks are better synchronized in Mcm23A mutants and genes in different cells are approximately at the same stage of their chromatin maturation process.

Slower forks are probably the reason why Mcm23A genes are captured at a later stage (i.e. step 2) of the chromatin maturation process than wt genes. The earliest stage of the process when all features are preferentially found on the strand that replicated first is not detected in Mcm23A cells. Instead, all genes in the mutant population are already on the next step: old nucleosomes are mostly bound to the strand that replicated first and new nucleosomes are mostly bound to its sister strand that replicated second. We propose that the slowing down of replication forks in the Mcm23A mutant changes the kinetics of chromatin maturation and favors the accumulation of chromatin configurations in step 2 of the process. This is supported by the fact that ΔRT distributions (the difference in replication timing between the lagging and the leading strand) for genes ordered by lagging/leading ratios of H3K56ac enrichment are inversely proportional to ΔRT distributions of genes that are ordered by lagging/leading ratios of H3K4me^3^ enrichment. In other words, H3K56ac is always enriched on the strand that replicated second and H3K4me^3^ is always enriched on the strand that replicated first (compare rows 1 and 2 with rows 3 and 4 from the top in **Figures 6 and S7 B**). In contrast to mutant cells, however, in wt cells in early-S phase both H3K56ac and H3K4me^3^ are enriched on the strand that replicated first (compare rows 1 and 3 and rows 1-4 in Figures 5 and S8A, respectively). Step 2 from mid-early S-phase was only captured in wt cells from replicate 1 (rows 2 and 4 **Figure 5**), when ΔRT distribution trends become inversely correlated with ΔRT trends from early S-phase, just like in Mcm23A cells.

We therefore conclude that chromatin maturation in Mcm23A mutants follows the same pattern as in wt cells. The observed differences between wt and mutant cells result from different kinetics of the same chromatin maturation process caused by the global slowing down of replication forks in the mutant.

### “Two-way” Two Step Model of Nucleosome Assembly and RNAPol2 binding to Daughter Genomes

The results presented above are consistent with the following model for chromatin maturation and RNAPol2 (re)binding on sister chromatids: a two-way two-step model of H3K4me3, H3K36me3 recycling, and H3K56ac binding followed by RNAPol2 binding and/or recycling to replicated gene copies (**Figure 7A**):

**Figure 7:**
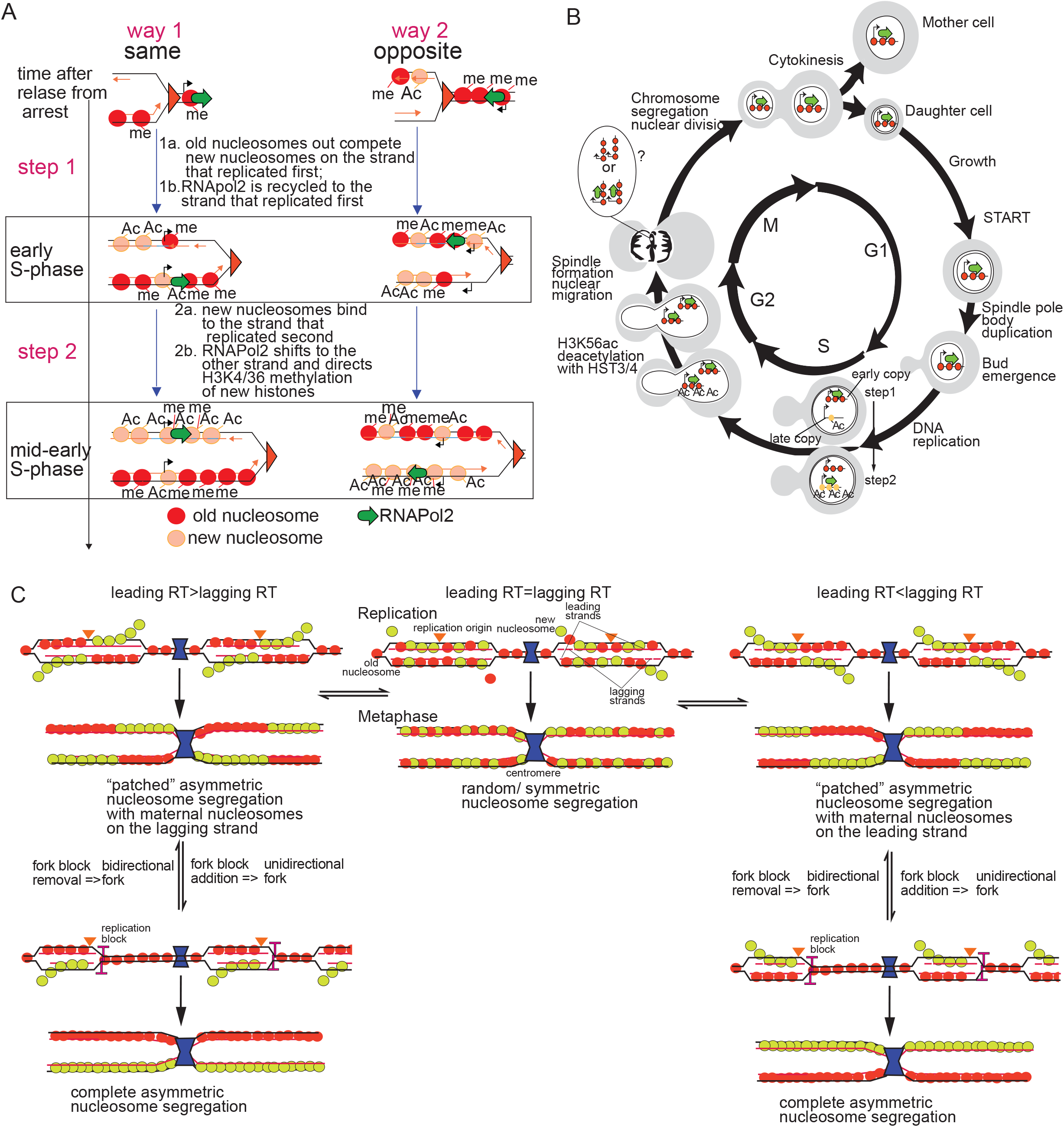
A. A “two-way” two-step model for chromatin assembly on daughter chromatids. Nucleosome deposition follows a two-step process. Step 1: “old” nucleosomes (red) and RNAPol2 (green arrow) binding first to the leading strand behind the fork while the lagging strand is still replicating when transcription and replication travel in the same direction (way 1). When transcription and replication travel in opposite directions, old nucleosomes are recycled on the lagging strand which replicated first (way 2). RNAPol2 will also bind to the lagging strand copy after the promoter has been replicated. New nucleosomes (tan) will be incorporated into the strand that replicated first mostly at promoters and ends of genes through replication independent turnover although some will outcompete old nucleosomes for binding to other sites in the CDS. Step 2: When replication of the other strand catches up it will be mostly populated by new nucleosomes. RNAPol2 also then apparently “switches” from the early replicating strand to the late one and directs H3K4 and H3K36 methylation of new histones by Set1 and Set2, respectively. **B.** Gene expression upregulation lags behind gene copy number doubling after replication. Since RNAPol2 is limiting immediately after replication only the earlier replicating gene copy containing old nucleosomes (red) is transcribed. Later on new acetylated nucleosomes (yellow) assemble on the second copy after it has finished replicating, and transcription shifts to that gene copy and directs the methylation of new nucleosomes. By G2, enough RNAPol2 molecules have been synthesized for both gene copies to be transcribed at pre replication levels. H3K56ac is removed at the G2/M transition. After deacetylation nucleosomes from the two daughter copies are indistinguishable and mother and daughter cells inherit identical chromatin configurations. **C.** Modulation of the replication timing of replicated gene copies determines the pattern of old and new nucleosome segregation.

#### Step 1

The two daughter chromatids are not replicated simultaneously at most genomic loci. Instead, one strand replicates a minute or more before the other. The leading strand usually replicates first at genes where transcription and replication go in the “same” direction (way 1). Conversely, the lagging strand replicates first at genes where transcription and replication go in “opposite” directions (way 2). Old nucleosomes (methylated on H3K4 and H3K36) are recycled behind the fork and initially compete with new nucleosomes for binding to the only replicated gene copy-the one that replicated first. Since the local concentration of old histones in the proximity of the replication fork is ostensibly higher than the local concentration of new histones, old nucleosomes out-compete new nucleosomes. RNAPol2 is at this stage also recycled to the strand that replicated first.

#### Step 2

New nucleosomes acetylated on H3K56 populate the sister strand that replicated later. RNAPol2 occupancy now shifts to the second strand as well, and directs methylation of H3 in new nucleosomes.

By mid-early S, the asymmetry in RNAPol2, H3K4/K36me^3^ and H3K56ac distributions is not detectable or appears diminished on genes that replicated the earliest because the signal comes from a mixture of step 1 and step 2 structures due to imperfect synchronicity of replication forks in different cells.

It is not yet clear why the replication machinery favors leading strand replication on “same” genes and “lagging” strand replication on “opposite” genes. Faster and earlier lagging replication on “opposite” genes is particularly puzzling. If the progress of the two DNA polymerases is coupled, as is currently thought, lagging strand synthesis should “naturally” lag behind leading strand synthesis, i.e. the same genome location should be replicated on the leading strand slightly before it is replicated on the lagging strand because the DNA strand that serves as a template for lagging strand synthesis has to fold into a loop to allow the leading and lagging strand DNA-polymerases to travel in the same direction(*24*). However, recent examination of the role of the Rad53 checkpoint kinase in replication stress uncovered that lagging strand synthesis outpaces leading strand synthesis when forks are stalled in the absence of Rad53(*25*). Leading and lagging strand replication can therefore be uncoupled under certain conditions. We propose that frequent encounters of the replication machinery with the transcription machinery, which are more likely to occur on “opposite” genes when the two complexes go towards each other, cause a progressive slowing down of the fork and transient uncoupling of leading and lagging strand synthesis.

Several observations support the hypothesis that forks become progressively slower when they go through a succession of “opposite” genes. First, the difference in replication timing between “opposite” and “same” genes increases with S-phase progression (**Figure S9A**). Second, genes of the same genic orientation tend to be replicated in succession by the same replication fork (**Figure S9B**). Consequently, a replication fork passing through an array of “opposite” genes is more likely to slow down compared to a fork replicating an array of “same” genes, presumably because of repeated successive and disruptive “head on” encounters with the transcription machinery in “opposite” gene clusters. Finally, leading strand synthesis is on average faster on same genes than on opposite genes, while lagging strand synthesis tends to be faster on opposite genes compared to same genes (**Figure S9C**). We hypothesize that the slowing down or transient pausing of the fork on opposite genes causes the temporary uncoupling of leading and lagging replication and favors lagging strand synthesis. Consequently, old nucleosomes and RNAPol2 are recycled on the lagging strand.

## Discussion

Our strand specific ChIP-NChAP technique enabled us to measure genome-wide binding dynamics of newly synthesized and old nucleosomes, and RNAPol2 complexes on two replicated daughter chromatids. Our results are consistent with a two-step “two-way” model of chromatin structure re-establishment after DNA replication that explains how the inherent asymmetry of old nucleosome and RNAPol2 distribution on replicated gene copies could either maintain or modulate gene expression states and chromatin configuration from one cell generation to the next (**Figure 7**).

It has recently been proposed that H3K56ac inhibits the expected upregulation of transcription caused by gene copy number doubling- a process that the authors of the study named transcription buffering (*19, 26*). Our results however do not support a direct role for H3K56ac in RNAPol2 binding to replicated genes. We propose instead that transcription buffering stems from the limiting local concentration of transcription factors (TFs) and RNAPol2 shortly after replication. The total transcriptional output from two gene copies after replication is still equal to the transcriptional output of one gene copy before replication simply because the locally available RNAPol2 pool is the same as in G1 (**Figure S1A**). Transcription resumes at “half capacity” shortly after replication using mostly locally available “recycled” TFs and RNAPol2 because it takes some time to synthesize sufficient amounts of new TFs and RNAPol2 complexes that are necessary for a two fold increase in transcriptional output. Consequently, transcription buffering, i.e. delaying until G2 the expected two-fold increase in gene expression, should last for a period of time that is needed to accumulate sufficient quantities of transcription machinery components.

Our analysis suggests that RNAPol2 is recycled behind the fork just as old nucleosomes are recycled. The distribution of “recycled” RNAPol2 complexes between daughter chromatids is not random and follows the same pattern as the distribution of old nucleosomes. We show that all nucleosomes and RNAPol2 are initially enriched on the strand that replicates first. Genic orientation seems to be the main factor that determines which strand will replicate first. Immediately after the passage of the fork old nucleosomes and some new nucleosomes are preferentially assembled on the early replicating strand because the other strand is not yet synthesized (step 1). The majority of new nucleosomes binds to the other strand after it has finished replicating because the strand that replicated first is by then already mostly occupied by old nucleosomes (step 2). RNAPol2 binding to replicated genes follows the same steps as nucleosome binding although with somewhat of a lag behind nucleosome assembly. Even though H3K56ac has a global stimulatory effect on transcription (Figure S5), the asymmetric binding and apparent switching of RNAPol2 from the copy that replicated first to the other copy, which is mostly occupied by new nucleosomes, is independent of rtt109 mediated H3K56 acetylation (Figure 4). RNAPol2 may still switch to the gene copy enriched for new nucleosomes due to their generally hyper acetylated state (*27–29*), although further experiments are needed to test this assumption. The transient increase in transcription shortly after replication that was observed in rtt109Δ cells (*19, 30*) is the basis for the hypothesis that H3K56ac is directly responsible for transcription buffering because it attenuates transcription of replicated genes during S-phase (*19*). Our results and the findings from Topal et al. (*30*), are however not consistent with this hypothesis. Indeed, Topal et al. (*30*) shows that gene expression goes back to a “buffered” state after an initial burst in transcription immediately after replication when H3K56ac is depleted. We therefore conclude that the observed transcription burst on replicated genes in rtt109Δ cells is caused by low nucleosome density on replicated DNA mmediately after replication due to the well documented defect in new nucleosome assembly in the absence of H3K56 acetylation (*31*) as suggested in Topal et al.(*30*).

According to our model, new histones should become methylated on H3K4me^3^, H3K36me^3^ and H3K79me^3^ when RNAPol2 switches from the gene copy that replicated first to its sister because the elongating RNAPol2 recruits Set1, Set2 and Dot1 methyltransferases, respectively. Finally, after global H3K56 deacetylation in late S (*32*) nucleosome configurations on the two sister copies become indistinguishable. By the start of the G2-phase, both copies carry the H3K4me^3^, H3K36me^3^ and H3K79me^3^ marks characteristic of transcribed genes. Consequently mother and daughter cells inherit gene copies with identical chromatin configurations (**Figure 7B**). The shift in RNAPol2 occupancy from one gene copy to the other thus ensures that both gene copies are transcribed in S-phase and that the same chromatin architecture is established on either copy even in conditions where RNAPol2 is limiting.

Our model based on relative differences in replication timing of the leading and lagging strands predicts that daughter chromatids will be “decorated” with contiguous alternating “patches” of old and new nucleosomes as illustrated in **Figure 7C** (left and right panels), as a consequence of the even distribution of replication origins along yeast chromosomes and the bi-directionality of replication forks. The other prediction of our model is that nearly simultaneous replication of both gene copies should reduce the bias recycling of old histones and result in a more random and symmetrical distribution as shown in the middle panel of **Figure 7C**. On the other hand, complete asymmetrical segregation of old and new histones could theoretically be achieved if replication fork barriers were introduced on the same side of all or most replication origins from the same chromosome and if the replication of the same strand happened consistently earlier than the replication of its sister strand throughout the chromosome (left and right panels, **Figure 7C**).

A recent study in murine ES cells proposed that new histones located in proximity of replication origins are slightly more enriched on the lagging strand (*16*), while a yeast study found a slight enrichment bias of new nucleosome on the leading strand (*14*). Our time course experiments spanning early to mid S-phase with parallel monitoring of the dynamics of five different chromatin features on thousands of gene copies from replicated sister chromatids in wt and mutant yeast cells now reveal a more complete picture. New histones are initially enriched either on the lagging or the leading strand depending on which strand replicated first. We propose that old nucleosomes outcompete new nucleosomes for binding to the only strand that is already replicated. At this stage both new and old nucleosomes are more abundant on that early replicating strand as the other strand has not been fully synthesized and has no nucleosomes on it (step1). New nucleosomes then bind to the sister strand after its replication is complete (step 2). Our model reconciles the findings of the studies above and suggests that they actually describe different steps and/or different types of loci that undergo the same chromatin maturation process.

Our finding that there is a substantial number of yeast genes with a new nucleosome bias for the leading strand in step 2 of the chromatin maturation process and the observation from Yu et al. (*14*) that new yeast nucleosomes have a slight “preference” for the leading strand are somewhat surprising given that the DNA polymerase processivity clamp PCNA is enriched on the lagging strand in all eukaryotes, including budding yeast ((*33*),reviewed in (*34*)). Since PCNA recruits the Chromatin Assembly Factor 1 (CAF1) to the replication fork (*35*) and CAF1 is responsible for the deposition of new H3-H4 tetramers on replicated DNA (*31, 36–39*), the expectation is that new histones will be enriched on the lagging copy. In light of our results and the findings from Yu et al. (*14*), the idea that the PCNA/CAF1 system is specialized for new nucleosome delivery to replicated DNA may need to be updated. It is possible that PCNA can also recruit a chaperone specific for old nucleosome re-assembly when the local concentration of old nucleosomes is higher than the concentration of new nucleosomes, which would enrich old nucleosomes on the lagging strand if the lagging strand is replicated first. Even though PCNA is enriched on the lagging strand, there are PCNA complexes in proximity of the fork on the leading strand as well. So, later on after leading strand replication has caught up, PCNA on the leading strand can recruit CAF1 bound to an Asf1/new H3-H4 complex and facilitate new nucleosome assembly on the leading strand. On the other hand, the Asf1 complex, which specializes in new nucleosome assembly, may not be the only partner for CAF1. It is possible that CAF1 might also interact with a chaperone complex that binds to old histones. This hypothetical CAF1/(old histone chaperone) complex could then also be targeted to nascent DNA through interactions with PCNA. The third possibility is that Asf1 also forms a complex with old nucleosomes and delivers them to CAF1. Indeed, it has recently been demonstrated that Asf1 is necessary for the recycling of old H3.3 and old H3.1 variants during transcription and replication in human cells, respectively (*40, 41*).

A nucleosome deposition system that is specialized for only one of the two replicated strands has been evoked for old nucleosomes as well. In order to explain a bias in old nucleosome deposition, three recent studies proposed a system of competing chaperone complexes that specialize in the preferential deposition of old histones on lagging or leading strands (*14–16*). According to this hypothesis, these competing nucleosome chaperons are Mcm2 and Dpb3/4 and they specialize in lagging and leading strand deposition, respectively. The Mcm2 subunit of the replication fork helicase Mcm2-7 has recently been implicated in the recycling of old nucleosomes behind the fork (*10*). Since it was observed that an Mcm2 mutation that impairs the interaction between histone H3 and Mcm2 apparently enhances the bias of old nucleosomes for the leading strand, two studies- one in yeast and one in mouse ES cells-have argued that Mcm2 promotes re-assembly of old nucleosomes specifically on the lagging strand. They proposed that Mcm2 mediated recycling of old nucleosomes to the lagging strand counteracts the “natural” tendency of old nucleosomes to re-bind to the leading strand (*15, 16*). Another yeast study has hypothesized that this “natural” predilection of old histones for the leading strand is actually orchestrated by the Dpb3/4 subunits of the leading strand DNA polymerase ɛ (*14*).

Models that are based on such highly specialized nucleosome deposition systems are however not entirely satisfactory. They don’t explain why one chaperone would consistently and ubiquitously outcompete the other. According to the conclusions of the above studies, Mcm2 in wt yeast seems to always “win” over Dpb3/4, resulting in a slight bias for old nucleosome deposition on the lagging strand while in mouse ES cells, Mcm2 consistently “loses” and old nucleosome bias leans more towards the leading strand. It is difficult to imagine a molecular mechanism based on specialized nucleosome chaperones that would create such opposing “natural” tendencies in two organisms that essentially have the same replication machinery.

Our experiments with the Mcm2 mutant show that old nucleosomes can be preferentially deposited on the leading or the lagging strand in equal proportions both in wt and Mcm23A cells alike. Mcm2 is therefore not likely to be a nucleosome chaperone that exclusively recycles old nucleosomes to the lagging strand since old nucleosomes are still recycled to the lagging strand in mutant Mcm23A cells. We therefore propose a model for nucleosome deposition that does not require specialized systems for old nucleosome deposition that are specific for the leading or the lagging strand. As described above, old nucleosome deposition depends instead on local replication dynamics. We propose that head on encounters of the transcription and replication complexes that are more likely to occur on “opposite” genes cause a transient uncoupling of leading and lagging strand synthesis, which in turn favors lagging replication. In that case old nucleosomes are deposited on the lagging strand first. Conversely, when replication and transcription travel in the same direction, the transcription machinery is less likely to impede the progress of the replication fork and leading and lagging strand synthesis remain coupled, thus favoring leading replication and consequently nucleosome deposition on the leading strand.

What is then the role of Mcm2 in old nucleosome recycling? As we have seen here, the Mcm23A mutant seems to only enhance the nucleosome deposition pattern already observed in wt cells. The major effect of the Mcm23A mutation appears to be instead a global slowing down of replication forks. We believe that this slowing down creates better synchrony between replication forks in different cells in the population. This synchrony of replication forks in the population produces a “more focused” snapshot of step 2 of the chromatin maturation process for three different chromatin features (H3K4me^3^, H3K36me^3^ and H3K56ac) on almost all replicated genes. Why does a mutation in Mcm2 that impairs the interaction between the helicase and histone H3 slow down replication forks? We hypothesize that the interaction between Mcm2 and H3 facilitates the removal of old nucleosomes ahead of the fork. If nucleosome removal is necessary for optimal DNA unwinding by the Mcm2-7 helicase, a defect in nucleosome removal would disrupt the optimal progression of the replication fork. So, when nucleosome removal is impaired, fork progression also slows down.

Along the same line of reasoning, we also suspect that the preferential deposition of old histones to the lagging strand observed in Dpb3/4 deletion mutants (*14*), is a consequence of a decrease in the rate of leading strand replication. We hypothesize that Dpb3/4 as subunits of the DNA polymerase ∊, probably stimulate the rate of leading strand synthesis. Consequently, leading strand synthesis should slow down in the absence of Dpb3/4 and old nucleosomes should be recycled to the lagging strand, which would have replicated first in these conditions.

We have seen so far that old nucleosome inheritance is asymmetrical with a bias for the leading or the lagging strand copy depending on which copy replicated first. This inheritance is for the most part accurate since the canonical distributions of H3K4me^3^ and H3K36me^3^ on gene bodies of recently replicated genes is largely preserved. In order for chromatin features to be truly epigenetic however, they also have to be instructive of the transcription state at their genomic location in addition to being accurately transmitted after cell division. We show here that the H3K4me^3^ and H3K36me^3^ marks that are recycled with old histones do not influence RNAPol2 distribution on replicated gene copies, despite having been accurately inherited. These marks found on actively transcribed genes are consequently not true epigenetic marks that convey information about active transcription to newly replicated gene copies and they probably do not direct the transcription machinery to continue transcribing the underlying replicated genes.

It has recently been shown that unlike these “active” histone marks above, gene “silencing” histone marks such as H3K27me^3^ and H3K9me^3^ in higher eukaryotes are inherited and instructive of the underlying silenced gene state (*42–44*). Thus, if such specific local chromatin configurations have to be inherited in only one daughter cell after cell division (either stem cell or differentiating cell for multicellular organisms) the bias in old nucleosome segregation at silenced loci could be enhanced by controlling the relative orientation of siRNA transcription and replication within heterochromatin. Opposing directionalities would favor deposition of “silencing” nucleosomes on the lagging strand and co-directionality would favor deposition on the leading strand.

Our model also provides a mechanistic blueprint for asymmetric nucleosome segregation that explains even the most extreme case of nucleosome segregation bias like the one recorded in Drosophila male germline stem cells (*45, 46*). There, the full complement of old nucleosomes is retained in the stem cell. As illustrated in Figure 7C, we speculate that such completely asymmetric segregation could be achieved with unidirectional replication forks as recent imaging data suggests (*47*) and transcription that mostly goes in the same direction throughout the chromosome.

Clearly, any locally asymmetrical nucleosome segregation would have to be coordinated with chromatid segregation during mitosis so that all the local chromatin configurations and gene expression states that are relevant for a specific cellular phenotype are (re)grouped in the same cell. One of the future challenges in understanding asymmetric cell divisions is therefore to decipher the molecular mechanism for coordinated co-segregation of genes with the relevant chromatin configurations and transcription states that are often dispersed on non-contiguous loci on one chromosome or even on entirely different chromosomes.

To summarize, we show that the recycling of chromatin features characteristic of actively transcribed chromatin is independent from the (re)binding of the transcription machinery to daughter chromatids. We suggest that the distribution pattern of both is constrained by the mechanics and interactions of replication and transcription ahead of the replication fork which determine which strand replicates first and then direct old nucleosomes and the transcription machinery to that strand. Our results suggest that RNAPol2 is also recycled behind the fork and is initially limiting, so only the gene copy that RNAPol2 landed on first is initially transcribed. The transcription machinery (while still limiting) switches to the sister strand later on where it directs the establishment of histone marks characteristic of active transcription onto new nucleosomes. Consequently, both replicated gene copies end up with the same chromatin features: one through inheritance of old histones and the other after the “inherited” RNAPol2 that binds there has recruited the relevant histone methyl-transferases.

So, the principal “epigenetic” factors that restore “active” chromatin features on most new histones are not old histones that serve as templates for a hypothetical copying mechanism that puts those marks on neighboring new histones. Our results indicate instead that the epigenetic factor that perpetuates active transcription states on newly replicated genes and their underlying “active” chromatin states is simply the recycled/inherited RNAPol2 that recruits histone methyltransferases that then deposit “active” marks on new histones.

## Materials and Methods

### Yeast Strains

All yeast strains have a W303 background and are listed in Table S3. The wt strain (RZ71) containing the HA tagged Rpb3 RNAPol2 subunit was obtained from a cross between ES3086 (courtesy of E. Schwob) and YMTK2567 (courtesy of T. Lee). The Rtt109Δ strain (RZ72) (Figure 5) was obtained by crossing RZ71 with ZGY929 (courtesy of Z. Zhang). The Rtt109Δ strain (RZ23) (Figure S3) was obtained from a cross between ES3086 and ZGY954 (courtesy of Z. Zhang). The Rtt109Δ (RZ72 and RZ23) and hst3, 4ΔΔ (PKY4220, courtesy of P. Kaufman, Figure S5). Strains have been additionally validated by anti H3K56ac western blotting of bulk mid-log cell extracts (data not shown). The Mcm2-3A strain (AC38) (Figures 6, S7, S8 and S9) was derived from a cross between strains 8566 (courtesy of K. Labib) and RZ71.

### Cell Culture

#### H3K56ac ChIP-NChAP (Figure 3)

Cells were grown overnight at 30°C in 500ml Synthetic Complete-URA + Dextrose (SCD-URA) media to OD 0.3. After 3.75hrs at 30°C with α factor (0.15μg/ml), cells were pelleted and transferred into preheated and premixed SCD-URA+ 10μM EdU (Carbosynth), with freshly added 20μg/ml pronase (Sigma). The culture was fixed with 1% formaldehyde after 24min (early S) or 30min (mid S) incubation at 30°C, incubated for 30min at 30°C and quenched with 125mM Glycine. Cell pellets were then washed with water and flash frozen in liquid nitrogen and kept at −80°C until further processing.

#### Rpb3-HA and H3K56ac time course ChIP-NChAP, synchronized (Figures 1, 2, 4, S1, S2, S3 and S4)

Cells were grown overnight at 30°C in SCD-URA. The culture was diluted to OD_600_ ~0.3 the next morning and grown to OD_600_ ~0.65 and re-diluted to OD_600_ ~0.3 (total final volume 10 L) in fresh media. The culture was synchronized with the addition of 0.15 μg/ml α factor for 3h30min at 30°C. Cells were released from arrest as above in preheated (SCD-URA) + 10 μM EdU. At 20, 22, 24, 25, or 32 min after release, cells (2.5 L per time point) were fixed with 1% (w/v) formaldehyde for 15 min at 30°C followed by 5 min of quenching in 125 mM Glycine. Cell pellets were then washed with cold PBS and flash frozen in liquid nitrogen and kept at −80°C until further processing.

#### Rpb3-HA, H3, H3K4me3, H3K36me3 and H3K56ac time course ChIP-NChAP, synchronized (Figures 5, 6, S6, and S7)

WT or Mcm2-3A cells were grown and synchronized as above. Cells from a 3 L culture were then fixed at 20 and 25 min after release from G1 arrest as above.

#### WT and Mcm23A NChAP time course for replication timing (Figure S8)

As above except that cells from 200 ml aliquots were fixed at 18, 25, 32, 40, 48 and 55 min after release from G1 arrest.

### MNase digestion

#### H3K56ac ChIP-NChAP (Figure 3)

700μl 0.5mm glass beads were added to frozen cell pellets (equivalent of 100ml cell cultures OD=0.5), re-suspended in 700μl cell breaking buffer (20% glycerol 100mM Tris-HCl 7.5). Cells were then spheroplasted by bead beating in the Bullet Blender (Next Advance) for 4×3min at strength 8 in the cold room. Spheroplasts were recovered by puncturing the cap of the tube and spinning into 5ml eppendorf at 1000rpm for 3 min. Cells were then centrifuged 5min at maximum speed in a micro centrifuge and the clear top layer was discarded, each pellet was re suspended in 600ul NP buffer (50mM NaCl, 10mM Tris-HCl pH 7.4, 5mM MgCl_2_, 1mM CaCl_2_,0.075% NP-40, 0.5mM sperimidine, 1mM βME). The amount of MNase (Worthington Biochemical) was adjusted to the cell density in each tube in order to obtain 80-90% mononucleosomal sized fragments after 20min incubation at 37°C. The reaction was stopped with 10mM EDTA and used for H3K56ac ChIP as described below.

### Chromatin Sonication

Cross-linked frozen cell pellets were re-suspended in 1.5 ml NP lysis buffer (100 mM NaCl, 10 mM Tris 7.4, 5 mM MgCl2, 1 mM CaCl2, 10% NP-40, 50 mM EDTA, 0.1 % SDS (optional), 1 mM PMSF and 1xEDTA-free protease inhibitor cocktail (Roche)). The suspension was then split into aliquots each containing ~10^9^ cells. Zirconium Sillicate beads (400 μl, 0.5 mm) were then added to each aliquot and cells were mechanically disrupted using a bullet blender (Next Advance) for 4 times x 3 min (intensity 8). Zirconium beads were removed from the cell lysate by centrifugation and the entire cell lysate was subject to sonication using the Bioruptor-Pico (Diagenode) for 3×10 cycles of 30 seconds *ON/*OFF each resulting in a final median size of chromatin fragments of 200 bp. Cellular debris was then removed by centrifugation and 2 % of the total supernatant volume was kept for the input and NChAP fractions.

### Chromatin Immunoprecipitation (ChIP)

#### H3K56ac (Figure 3)

All steps were done at 4°C unless otherwise indicated. For each aliquot, Buffer L (50 mM Hepes-KOH pH 7.5, 140 mM NaCl, 1 mM EDTA, 1% Triton X-100, 0.1% sodium deoxycholate) components were added from concentrated stocks (10-20X) for a total volume of 0.8 ml per aliquot. Each aliquot was rotated for 1 hour with 100 μl 50% Sepharose Protein A Fast-Flow bead slurry (IPA400HC, Repligen) previously equilibrated in Buffer L. The beads were pelleted at 3000 X g for 30sec, and approximately 100 μl of the supernatant was set aside for the input sample. With the remainder, antibodies were added to each aliquot (equivalent to 100 ml of cell culture): 6μl anti- H3K56ac for the mid-S time point (Merck-Millipore, 07-677-IS (lot# 266732) or 10 μl anti- H3K56ac for the early-S time point (Active Motif, 39281 (lot# 14013003). Immunoprecipitation, washing, protein degradation, and DNA isolation were performed as previously described (*48*). Purified DNA was treated with RNAse A (Qiagen) and CIP (NEB) and purified once more with Phenol-Chloroform. Fragments shorter than 100bp were removed with homemade MagNA beads (SeraMag Speed beads, Thermo Scientific,(*49*)), and purified fragments were used for NGS library construction (Input, ChIP) or biotin conjugation and subsequent NGS library construction (NChAP, ChIP-NChAP).

#### Rpb3-HA, H3, H3K4me3, H3K36me3 and H3K56ac (Figs. 1, 2, 4, 5, 6, S1, S2, S3, S4, S6, S7)

Sonicated chromatin was precleared using Protein A agarose beads (Repligen) for 1 hour (1h) at 4°C on the rotating wheel. The sonicated material was then pooled together and distributed into 500 μL aliquots (equivalent of 7*10^8^ cells per aliquot) and 25 μl of Protein G magnetic beads (Life Technologies-Invitrogen) pre-bound with 6μg, 3μg, 2μg, 4μg, or 3μg of anti-HA (ab9110, abcam) or anti-H3K56ac (Active motif, 39281), anti H3K4me3 (abcam, ab8580), anti H3K36me3 (abcam, ab9050) or anti-H3 (abcam, ab1791) respectively, was added to each tube. Aliquots were then incubated with rotation at 4°C overnight. The beads were then washed once with cold buffer L (50 mM HEPES-KOH, pH 7.5, 140 mM NaCl, 1 mM EDTA, 1% Triton X-100, 0.1% Na-deoxycholate), three times with cold Buffer W1 (Buffer L with 500mM NaCl), twice with cold Buffer W2 (10 mM Tris-HCl, pH 8.0, 250 mM LiCl, 0.5% NP-40, 0.5% sodium deoxycholate, 1 mM EDTA), and once with cold TE buffer (10 mM Tris-HCl,pH 8.0, 1 mM EDTA). Chromatin was eluted in 2×125 μl elution buffer (25 mM Tris-HCl, pH 7.5, 5 mM EDTA, 0.5 % SDS) by incubation for 20 min at 65°C. The eluates and reserved input samples were treated with RNase A (Qiagen) for 1h in 37°C and proteins were then digested with Proteinase K (Euromedex, final concentration 0.4 mg/ml) for 2h at 37°C and the temperature was then shifted to 65°C overnight to reverse cross-links. DNA was then purified with the QIAquick PCR purification kit (QIAGEN) except for the early S-phase time point (54% non-replicated) from Figs. 1B and 2 that was purified by Phenol Chloroform extraction to keep fragments smaller than 100 bp, which increases the resolution for mapping replication origins. The specificity of H3K56ac antibodies was tested by Western Blot using an H3K56A mutant strain. (Figure S10).

### Biotin conjugation to EdU with the Click reaction

10μl DNA solution was mixed with 10μl biotin azide (quanta biosystems) solution in DMSO/tBuOH(3:1). For each pmole of DNA, we added 1mM biotin azide solution (for example for 20pmoles of DNA in 10μl, we added 10μl 20mM biotin azide). 10μl CuBr solution (10mM CuBr (from freshly made stock), 10mM TBTA (Eurogentec), 10mM Ascorbic acid (from freshly made stock) in DMSO/tBuOH 3:1) were then added to the DNA-biotin azide mix and the reaction was shaken for 2hrs at 37°C. 300μl 10mMTris-HCl pH7.5, 8μl 0.25% linearized acrylamide solution, 33μl 3M Sodium Acetate pH5 and 1ml 100% cold EtOH were then added to the Click reaction and DNA was precipitated at −20°C overnight.

### Illumina Sequencing Library Construction

#### ChIP-NChAP and NChAP libraries (Figures 1B, 2, 3, 4, S2B, S3 and S4)

Biotinylated DNA pellets were re suspended in 25μl TNE0.2 buffer (200mM NaCl, 10mMTris-HCl 7.5, 1mM EDTA) and mixed with 25μl Streptavidin coated magnetic beads (NEB, pre washed in TNE0.2 and blocked with 100μg/ml salmon sperm DNA). The DNA and bead mixture was shaken for 30min at RT, and beads were washed 2x with 0.25ml TNE0.2 buffer and re suspend in 35μl 10mM Tris-HCl pH8. All the subsequent steps were done with DNA attached to the beads. DNA fragments were blunt ended and phosphorylated with the Epicentre End-it-Repair kit (1X buffer, 0.25mM dNTPs,1mM ATP, 1μl Enzyme mix in a 50μl reaction) for 1hr at RT. Beads were washed 2x with TNE0.2 and re suspended in 43μl 10mM Tris-HCl pH8. Adenosine nucleotide overhangs were added using Epicentre exo- Klenow for 45min at RT (with 0.2mM dATP) followed by two TNE0.2 washes and re suspension in 15μl 10mM Tris-HCl pH8. Illumina Genome sequencing adaptors with in line barcodes (PE1-NNNNN: PhosNNNNNAGATCGGAAGAGCGGTTCAGCAGGAATGCCGAG PE2-NNNNN: ACACTCTTTCCCTACACGACGCTCTTCCGATCTNNNNNT, NNNNN indicates the position of the 5bp barcode, (IDT)) were then ligated over night at 16°C using the Epicentre Fast-Link ligation kit. The ligation reaction was washed 2x with TNE0.2 and beads were re suspended in 20μl water. DNA was then subjected to a primer extension reaction with dUTP to separate the nascent strand from its complement (1X NEB buffer2, 0.1μg/μl 5’phosphorylated random hexamers (IDT), 1.72 μM Illumina PE primer 2.0 (IDT), 0.6 mM dNTPs (dUTP instead of dTTP) and 2U/μl Klenow 5NEB). DNA was denatured and annealed to the primers prior to enzyme addition and the reaction was incubated 1.5 hrs at 37°C. Beads were washed 4x and re suspended in 20μL water. The dUTP containing strand was degraded with USER enzyme (NEB) and beads were re suspended after washing in 20μl 10mM Tris-HCl pH8. The remaining nascent DNA strand was amplified with the Phusion enzyme (NEB) for 18 PCR cycles with Illumina PE1 (AATGATACGGCGACCACCGAGATCTACACTCTTTCCCTACACGACGCTCTTCCGATCT) and PE2 (CAAGCAGAAGACGGCATACGAGATCGGTCTCGGCATTCCTGCTGAACCGCTCTTCCGATCT) primers (IDT). Only 2μl of the bead suspension was added to the 50μl PCR mix. Amplified libraries were purified using MagNA beads (SeraMag Speed beads, Thermo Scientific,(*49*)) and fragment size and library concentration were determined from a Bioanalyzer (Agilent) scan and Qubit fluorimetry measurements, respectively.

#### ChIP-NChAP and NChAP libraries (Figures 5, 6, S6, S7,and S8)

As above except the TrueSeq V2 LT Sample prep kit (Illumina) was used for the blunt-ending, A tailing and adaptor ligation steps. The primer mix from the kit was also used in the PCR amplification step (15 cycles).

#### Input and ChIP libraries

Libraries were constructed as above from the blunt ending step. DNA fragments were blunt ended and phosphorylated with the Epicentre End-it-Repair kit. Adenosine nucleotide overhangs were added using Epicentre exo- Klenow. Illumina Genome sequencing adaptors with in line barcodes (above) were then ligated over night at 16°C using the Epicentre Fast-Link ligation kit. Ligated fragments were amplified as above using the Phusion enzyme. Reactions were cleaned between each step using MagNa beads. Libraries for input and ChIP (H3K56ac and Rpb3-HA) fractions from replicates 1 and 2 (52%, 45% and 38% non-replicated) from Fig. S2, for input and Rpb3-HA ChIP from rtt109Δ replicates (17% and 10% non-replicated) from Fig. 4, and for input and all ChIPs in Figures 5, 6, and S7 were prepared using the TrueSeq V2 LT Sample prep kit (Illumina). The libraries of input fractions from Figure S8 were prepared using the NEBNext^®^ Ultra™ II DNA Library Prep Kit for illumina (NEB).

### Illumina Sequencing

Libraries were mixed in equimolar amounts (10 to 15 libraries per pool) and library pools were sequenced on a HiSeq 2000 (2×75bp) (Illumina) at the CNAG, Barcelona, Spain or the NextSeq550 (2×75bp) (Plateforme Transcriptome, IRMB, Montpellier, France).

### RNAseq with spike-in control (Figure S5)

Exponentially growing S.cerevisiae and S. pombe (strain FY2319, courtesy of S. Forsburg) cells were flash frozen in liquid N2 and total RNA was isolated from frozen cell pellets with Trizol. Frozen cell pellets were re-suspended directly in Trizol and bead beated in the Bullet Blender (Next Advance) as above. RNA was then purified and DNAseI treated with the RNAeasy Column purification kit (Qiagen). Extracted total RNA amounts were measured on the Qubit and the Nanodrop and the quality was checked in a Bioanalyzer scan (Agilent). Each S.cerevisiae total RNA extract was mixed with the S.pombe total RNA extract at a mass ratio of 10:1. The mixed RNA samples were then used for NGS library preparation using the Illumina TruSeq Stranded mRNA kit according to the manufacturer’s protocol.

### ChIP DNA Microarray hybridization (Figure S1A)

ChIPed DNA and their corresponding input samples were amplified, with a starting amount of up to 30 ng, using the DNA linear amplification method described previously (*48, 50*). 2.5 μg of aRNA from each sample produced from the linear amplification was transformed into cDNA by reverse transcription in the presence of amino-allyl dUTP. The resulting cDNA was dye-coupled with Cy5 or Cy3 NHS-esters and purified as described previously (*50*).

Labeled probes (a mixture of Cy5 labeled input and Cy3 labeled ChIPed material or their corresponding dye flips) were hybridized onto an Agilent yeast 4×44 whole genome array. Images were scanned at 5μm with the InnoScan 710 MicroArray scanner (Innopsys) and processed with the Mapix software. Data was normalized by dividing the Cy3/Cy5 (or Cy5/Cy3 ratio for the dye flip) ratio for each probe with the average Cy3/Cy5 ratio for the whole array. The average of the pair of normalized ratios from the dye flip technical replicates was used in the final analysis. The GEO accession number for the microarray data is

### Western Blot (Figure S10)

10 mL aliquots from time points above were mixed with 2 mL 100% TCA and kept on ice for 10 min. Cells were then pelleted and washed twice with 500 μL 10% cold TCA. Pellets were resuspended in 300 μL 10% cold TCA and bead beated with Zirconium Sillicate beads (0.5 mm) in a bullet blender (Next Advance) for 3 times x 3 min (intensity 8). Zirconium beads were removed from the cell lysate by centrifugation and the entire cell lysate was washed twice with 200 μL 10% cold TCA. The cells lysate was then pelleted and re-suspended in 70 μL 2xSDS loading buffer (125 mM Tris pH 6.8, 20% glycerol; 4% SDS, 10% β-mercaptoethanol, 0,004% bromophenol blue) preheated at 95°C. Approximately 30 μL Tris (1M, pH 8,7) was added to each sample to stabilize the pH. Samples were heated for 10 min at 95°C, pelleted and the soluble protein extract in the supernatant was transferred to new tubes. Protein concentrations were measured by Bradford test kit (Sigma, B6916) and 10 μg/sample was loaded on a 15% polyacrylamide SDS-PAGE gel (30:1 acrylamide/bis-acrylamide). Proteins were transferred after electrophoresis to a PDVF membrane (Bio-Rad, 1620177). The membrane was incubated for 1h at room temperature with anti-H3K56ac (Active motif, 39281) at 1:2000 dilution. Secondary goat anti-rabbit-HRP (1:10000, Santa Cruz Biotechnology sc-2054) was added after the primary antibody and incubated for 1h at room temperature. All antibodies (primary and secondary) were diluted in 5% milk/TBS. The membrane was washed 3x in 1xTBS-10%Tween after each antibody incubation step. The blot was then covered with 500 μL Immobilon Forte Western HRP substrate (Millipore WBLUF0500) for 2 min and protein bands were detected on a high-performance chemiluminescent film (Amersham 28906837).

### Data Analysis

All analysis was done using in house Perl and R scripts available upon request.

#### ChIP, ChIP-NChAP, NChAP

Sequences were aligned to S. Cerevisiae genome using BLAT (Kent Informatics, http://hgdownload.soe.ucsc.edu/admin/). We kept reads that had at least one uniquely aligned 100% match in the paired end pair. Read count distribution was determined in 1bp windows and then normalized to 1 by dividing each base pair count with the genome-wide average base-pair count. Forward and reverse reads were treated separately.

The repetitive regions map was constructed by “BLATing” all the possible 70 bp sequences of the yeast genome and parsing all the unique 70bp sequences. All the base coordinates that were not in those unique sequences were considered repetitive.

Normalized read densities for all genes were aligned by the transcription start site (*51*) and median read densities for each coding region (from the tss to the transcription termination site) were determined for all datasets. Median read densities from ChIP and NChAP (nascent chromatin) fractions were normalized to the median from their corresponding input (sonicated or MNAse digested chromatin) and medians from ChIP-NChAP fractions were then normalized to the corresponding input normalized ChIP fraction.

#### Replicated genome fraction

Normalized read counts, binned in 400bp windows over the whole genome, from NChAP fractions (and the H3K56ac mid S ChIP fraction, Fig. 3) in each chromosome were divided by the maximum read count for that chromosome to obtain population read densities (i.e. the fraction of the cell population in which each 400bp genome segment has been replicated). We then determined the distribution of these read densities into 100 bins from 1% to 100%. The non-replicated fraction was the genome fraction with read densities between 0 and 1%.

#### Replication origins mapping

Origins were mapped from the nascent chromatin fraction in the early S-phase dataset from Figure 1C and Figure 2 (54% non-replicated). The resolution for origin centers was higher in this dataset because small fragments (<100bp) were not removed from this fraction (see the Chromatin Immunoprecipitation (ChIP) section). We identified local peaks within Replication Origin Associated Domains (ROADS, i.e. replicated regions around known origins of replication) on every chromosome (Table S1). We then looked for ACS consensus sequences (*52*) within +−200bp of each identified peak and kept the ACS sequence closest to the peak (Table S2). Peaks without ACS sequences were eliminated from further analysis.

#### Replication timing (Figure S8 A-E)

Replication timing in Figure S8A was determined as described in (*22*). Watson and Crick reads from each time point were first normalized to input and then normalized to 1 per chromosome by dividing each 50bp segment density with the highest density in the chromosome. The normalized densities were then smoothed with a 1600bp moving window average. Read densities for each 50bp segment were then plotted against time. Replication timing was determined from the Hill equation fit: 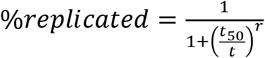, where *t* is the time since release from arrest and the *t_50_* is replication timing, i.e. the time since release at which that 50bp segment has been replicated in 50% of the population.

Replication timing values per gene copy are listed in Table S4.

#### DNA synthesis rates (Figure S6 B and Figure S8 F-H)

First, replication origins were determined as above from NChAP fractions of 18 et 25min time points (Figure S8) or 20 and 25min time points (Figure S6B). Local peaks were found in read density profiles binned in 50bp windows and normalized as above for replication timing. Second, genes that are replicated from each origin were grouped into Replication Origin Associated Domains (ROADs) from Watson and Crick strands in both time points. The average DNA synthesis rates of the Watson and Crick strands at each gene from every ROAD were calculated from the slope of the linear fit of the gene coordinates in bp versus replication timing. Synthesis rates of lagging strand gene copies are derived from slopes from the Watson strand upstream of origins and the Crick strand downstream of origins. Conversely leading strand synthesis rates come from Crick strands upstream and Watson strands downstream of origins.

DNA synthesis rates per gene copy with ROADs from Figure S6A are listed in Table S5.

#### RNA-seq normalized to S.Pombe spike-in (Figure S5)

S.Cerevisiae and S.pombe reads were aligned to their respective genomes using BLAT and the read density distribution was determined for each species in each dataset separately. The average S.pombe genomic read density per bp (F and R reads were processed together) was determined for each dataset. For spike-in normalization, S.Cerevisiae read densities per bp were then divided with the corresponding S.pombe average genomic read density. For internal normalization S.Cerevisiae read densities were divided with its average genomic read density as described above. Normalized read densities for each gene were aligned by the transcription start site and divided into sense and antisense transcripts. The median read density for each gene (from the tss to the end of the coding sequence) was then determined for each transcript. Intron regions were excluded from the calculation.

## Author Contributions

RZ optimized ChIP-NChAP and performed all the experiments except the ones in Figure 3 (done by MRL) and Figures S5 and S10 (done by AC). AC assisted RZ in the cell culture and ChIP steps of the experiments. RZ and AC constructed the yeast strains. MRL, RZ and AL designed the experiments. MRL conceived and developed ChIP-NChAP, wrote the Perl and R analysis scripts, analyzed the data, and wrote the manuscript.

## Acknowledgments

We thank Susan Forsburg, Paul Kaufman, Traci Lee, Etienne Schwob, Karim Labib and Zhiguo Zhang for yeast strains. Thank you to Marta Gut and Julie Blanc from CNAG (Barcelona, Spain) and Veronique Pantesco (Plateforme Transcriptome-IRMB Montpellier) for deep-sequencing services. Thank you to Robert Feil for critical reading of the manuscript. This work was supported with the ERC-Consolidator grant- NChIP 647618 (MRL).

