## Supplementary Figures for "Mechanics of DNA Replication and Transcription Guide the Asymmetric Distribution of RNAPol2 and Nucleosomes on Replicated Daughter Genomes"

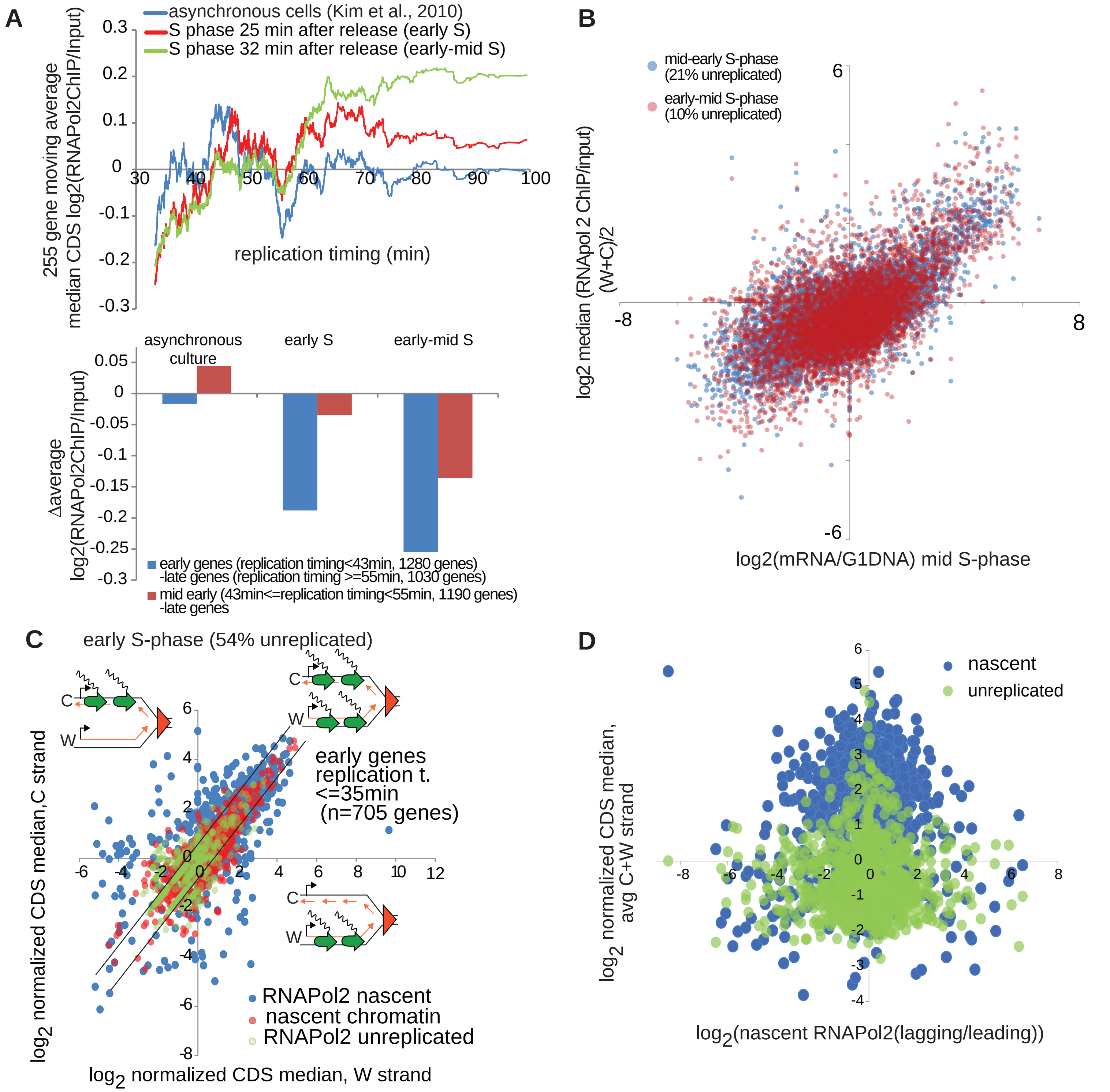


**Figure S1: RNAPol2 is limiting after replication and is distributed more asymmetrically on replicated chromatids at genes with low RNAPol2 density. A.** Correlation between the 255 gene moving window average of the median RNAPol2 occupancy (RNAPol2 ChIP/input) in the coding sequence of each yeast gene (excluding promoters) and gene replication timing (Vasseur et al., 2016). RNA pol2 occupancy was measured in synchronized wt cells in early (25min after release from G1 arrest) and early-mid (32min) S-phase by HA tagged RNAPol2 ChIP hybridized to whole genome two channel microarrays (4x44K Agilent), occupancy values are an average of two dye flip technical replicates (Top). Bottom: Difference in average log2(RNAPol2/input) between early and late genes (blue) and mid early and late genes (red) in asynchronous cultures, and in early and mid-early S-phase. As replication in the cell population progresses RNAPol2 occupancy relative to gene copy number decreases, i.e. earlier replicated genes have less RNAPol2 per gene copy than late genes that have not yet been replicated; compare early replicating genes (replication timing <43min) or mid early replicating genes (43min<=replication timing<55min) to late replicating genes (replication timing >=55min) in red (early S) and green (mid early S) curves in the top panel, and in the bar graph in the bottom panel. Conversely RNAPol2 occupancy in asynchronous cells (blue, (52)) shows the expected pattern of higher occupancy in early genes compared to late genes as early genes are known to have on average higher transcriptional activity than late replicating genes. Ratio values on the Y axis have been normalized to 0 by subtracting the average log2(RNAPol2/input) for all genes from the log2(RNAPol2/input) for each gene. Release media contained 10μM EdU. **B.** Correlation between bulk RNAPol2 occupancies in mid-early and early-mid S (repl. 1) and mRNA abundance in mid S. (from heat map in Figure 2A). **C.** W versus C copies scatter plot of median normalized (as in Figire 2A) read densities of early replicating genes(shown in Figure 2A) for RNAPol2 on nascent chromatin (ChIP-NChAP fraction, blue), nascent chromatin (NChAP fraction, red), and RNAPol2 on unreplicated chromatin (ChIP unreplicated fraction, green). The biggest differences between the two replicated copies are seen in the ChIP-NChAP fraction, suggesting asymmetrical distribution of RNAPol2. **D.** Scatter plot of the ratio of RNAPol2 occupancy between the lagging and the leading gene copy for all 705 early genes from C and the average median RNAPol2 density (average of W and C copies) on nascent chromatin (blue) and unreplicated chromatin (green). Genes with the largest difference in lagging and leading RNAPol2 occupancy after replication have low expression and low RNAPol2 density .

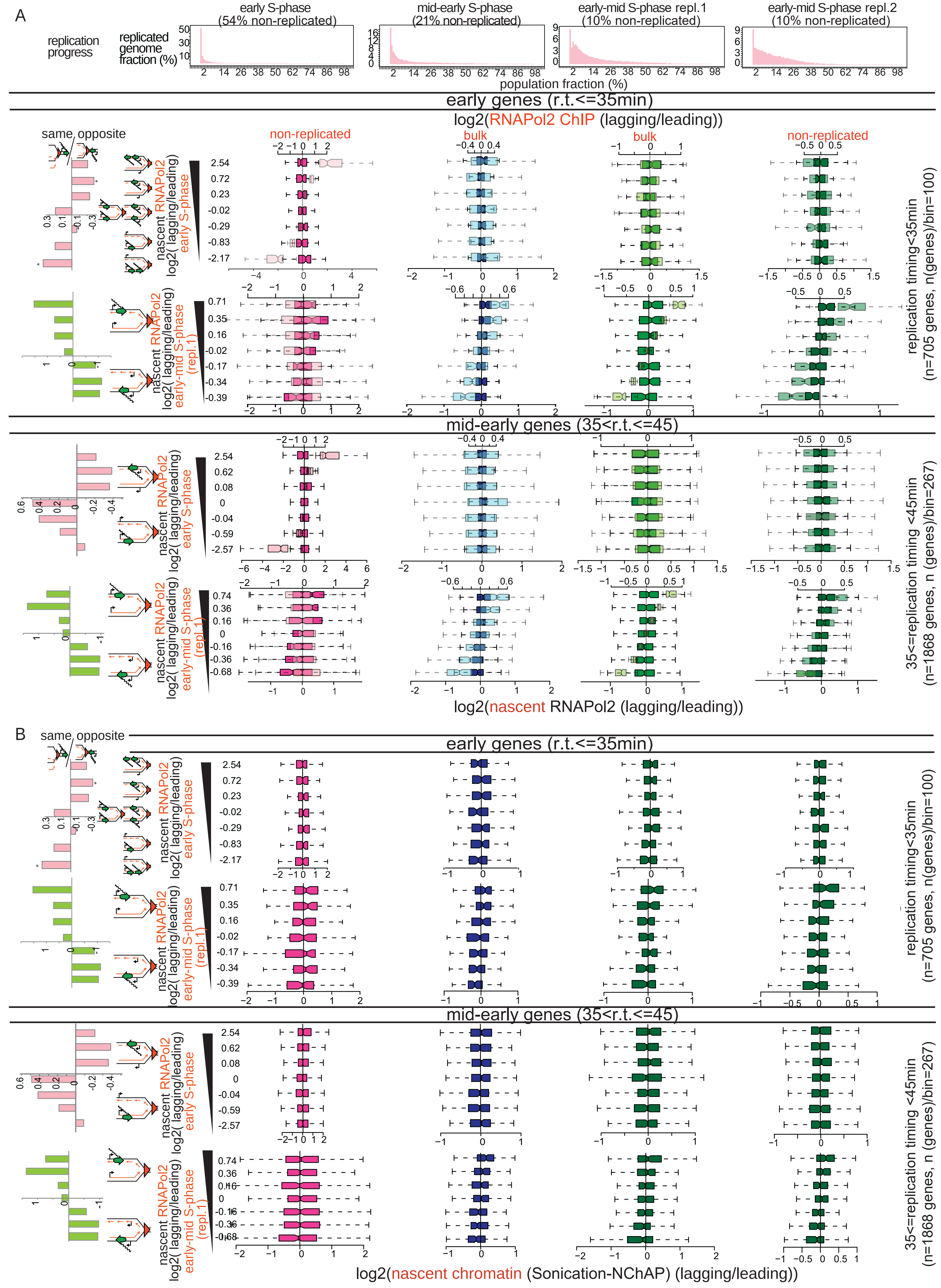


**Figure S2: Lagging and Leading strand read density distributions from ChIP (bulk and non-replicated) and NChAP (nascent chromatin) are symmetrical.** A As in Figure 2B with log2 (RNAPol2 ChIP (lagging/leading)) box plots in dark superimposed on log 2(nascent RNAPol2-ChIP-NChAP(lagging/leading)) distributions in light boxes. B log2 (nascent chromatin (NChAP)(lagging/leading)) box plot distributions.

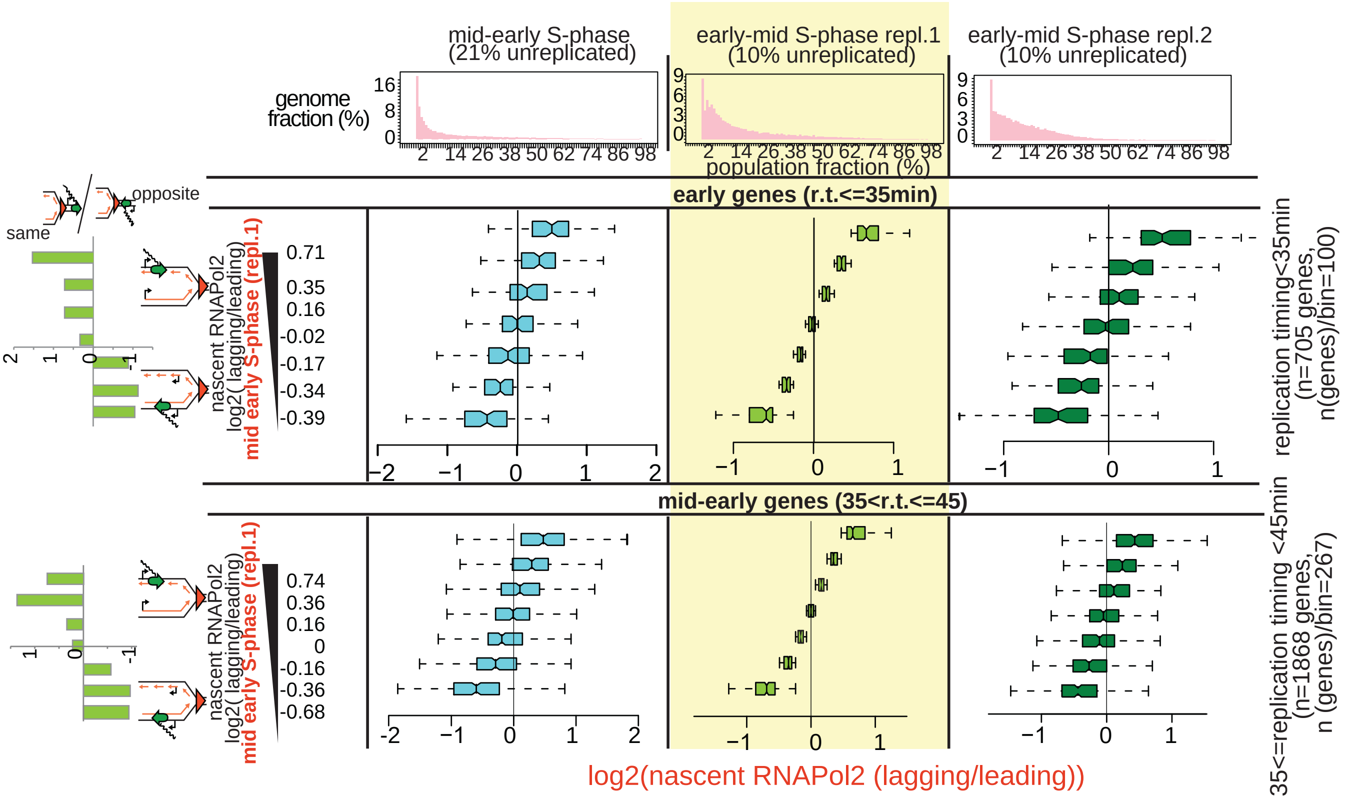


**Figure S3: The RNAPol2 distribution pattern from early-mid S-phase is reproducible.** Box plot distributions of lagging/leading nascent RNAPol2 ratios from mid-early and early-mid S-phase (from Figure 2A) (columns left to right) for early and mid-early genes (1st and 2nd row from the top, respectively). The header row shows the distribution of genome read densities (in 400bp bins) normalized to the maximum read density for each NChAP fraction (reads have not been normalized to input) at indicated time points in S-phase. Early and mid-early genes in rows 1 and 2, respectively have been sorted by decreasing lagging/leading RNAPol2 occupancy in early-mid S-phase (replicate 1, middle column, yellow background) and divided into 7 bins (y axis), and box plot distribution of nascent RNAPol2 lagging/leading ratios (x axis) have been determined for each bin in the replicates (columns 1 and 3). For example, early genes (705 genes shown in Figure 2A) were sorted by increasing lagging/leading nascent RNAPol2 ratio from the early-mid S-phase (replicate 1) and then divided into 7 bins of ~100 genes each. We determined the box plot distribution of lagging/leading ratios for each bin in each replicate. The average lagging/leading ratios from the early-mid S replicate 1 dataset for each bin are in the y axis on the left. For example the bottom group of genes in row 1 has on average 30% more RNAPol2 on the leading copy than on the lagging in early-mid S-phase. We then calculated the ratio of “same” orientation genes versus “opposite” genes for each group normalized to the same/opposite ratio of all 705 early genes (row 1) or 1868 mid early genes (row 2) (bar graphs, on the left). The bar graphs show the “same” gene enrichment for gene bins indicated in the Y axis of each row on the right. “Same” gene enrichment is proportional to the nascent RNAPol2 lagging/leading ratio, i.e. “same” genes in early-mid S-phase tend to have more RNAPol2 on the lagging copy and “opposite” genes tend to have more RNAPol2 on the leading copy. The same pattern of RNAPol2 distribution is detected in mid-early S-phase (column 1). RNAPol2 lagging/leading ratios from a replicate time-point in early-mid S-phase (replicate 2, column 3) also correlate with replicate 1.

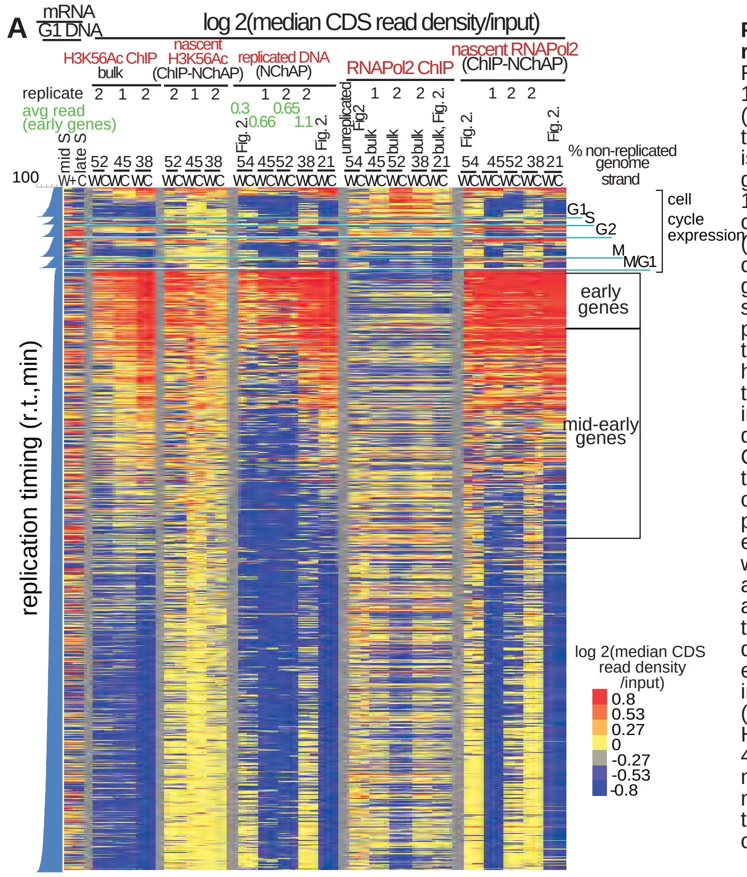


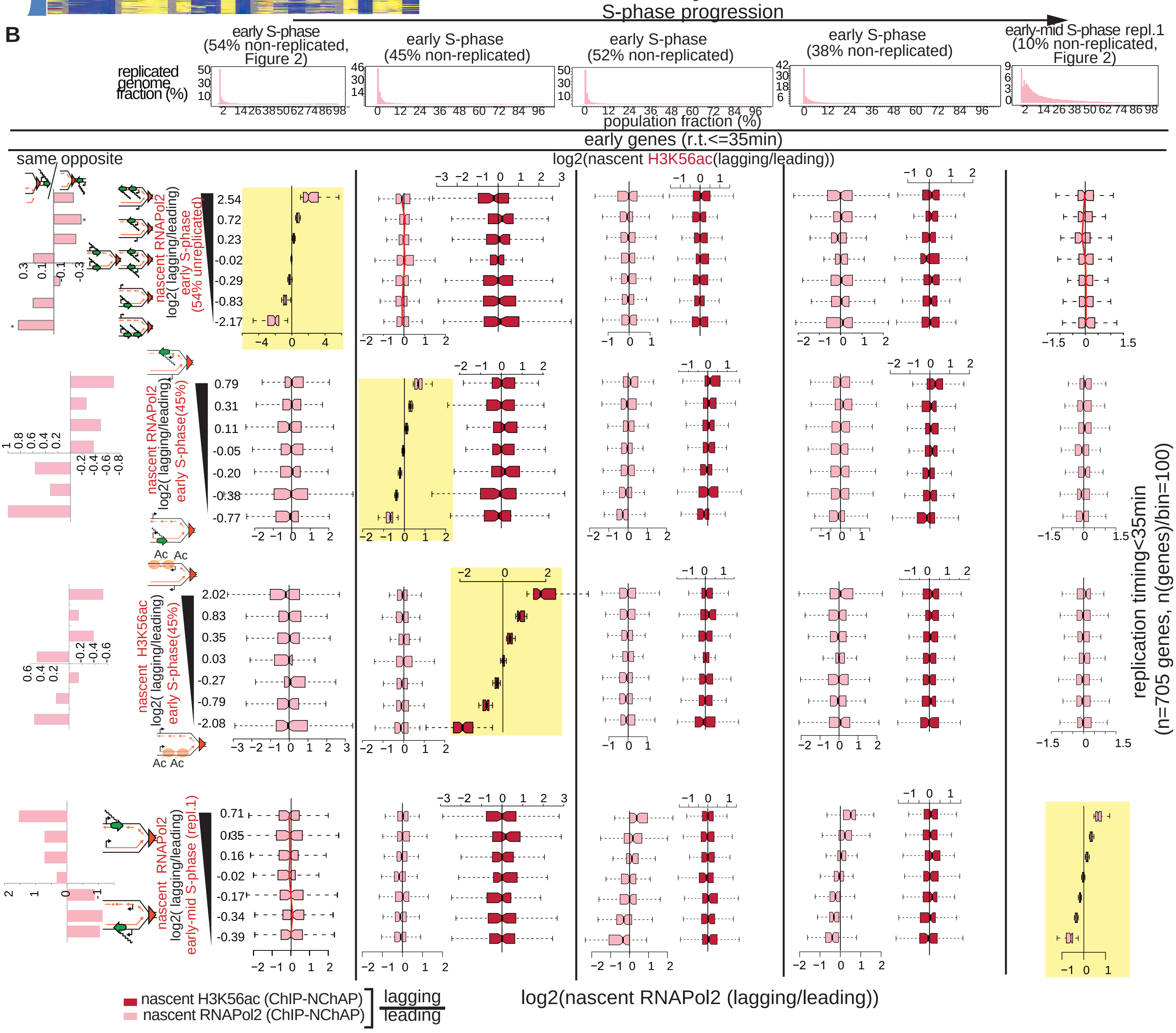


**Figure S4: Asymmetric distribution patterns of RNAPol2 and H3K56Ac on replicated gene copies do not correlate:replicate time-course in early S-phase. A.** Replicate time course experiments in early S-phase were performed as outlined in Figure 1. The Heat map shows median RNAPol2 and H3K56Ac occupancies in coding regions (CDS) of all yeast genes. H3K56Ac and RNAPol2 ChIPs were performed in parallel from the same cell culture as indicated by the replicate number above the heat map. Each line is an individual gene and columns represent occupancy values for (W)atson and (C)rick gene copies for different S-phase time points : early S (45% non-replicated (time course 1), 52% and 38% replicated (time course 2) ) from the ChIP, NChAP (sonicated nascent chromatin) and ChIP-NChAP (Nascent RNAPol2 and H3K56Ac) fractions. Early-mid S (21% non-replicated) and early (54% non-replicated) datasets from Fig. 2 are added for comparison. The first two columns on the left represent mRNA enrichment over G1 genomic DNA in mid and late S (in the absence of EdU) determined with gene expression microarrays (Vasseur et al., 2016). Genes are grouped by cell cycle expression patterns and then ordered by replication timing within each group as in Fig. 2(replication timing determined in Vasseur et al., 2016). Median read density values for each gene have been normalized by separately dividing the W and C read densities of each gene in the ChIP and NChAP fractions with the W and C average read density of the sonicated input fraction for the same gene. The ChIP-NChAP medians have been normalized by dividing the W and C values with the input normalized W and C average values from the ChIP fraction. Due to the stochastic nature of replication origin activation it is impossible to precisely reproduce each early S-phase time point from one biological replicate to the other. Early S-phase time points were therefore further sorted according to replication progression by calculating the average median NChAP density of W and C reads for early genes (n=705), which shows that the replicate from Figure 2 (54% non-replicated) with an average NChAP read density of 0.3 is the earliest time point, followed by the 45% and the 52% non-replicated points, which are at the same stage in S-phase (with 0.66 and 0.65 average read densities, respectively) and finally with the 38% non-replicated time-point as the latest in early S (1.1 average read density of early genes). **B.** Box plot distributions of lagging/leading nascent RNA pol2 and H3K56Ac ratios from early to early-mid S-phase (columns left to right) for early genes. Genes have been organized into bins of increasing lagging/reading ratios (yellow background) for RNAPol2 in early S ( dataset from Figure 2; row 1), early S (dataset from A, 45% unreplicated; row2), H3K56ac in early S (45% unreplicated; row 3) and early-mid S (Figure 2, replicate 1; row 4). The header row shows the distribution of genome read densities (in 400bp bins) normalized to the maximum read density for each NChAP fraction (reads have not been normalized to input) at indicated time points in S-phase. The bar graphs on the left show the "same" gene enrichment calculated as in Fig. 2C for gene bins indicated in the Y axis of each row on the right.

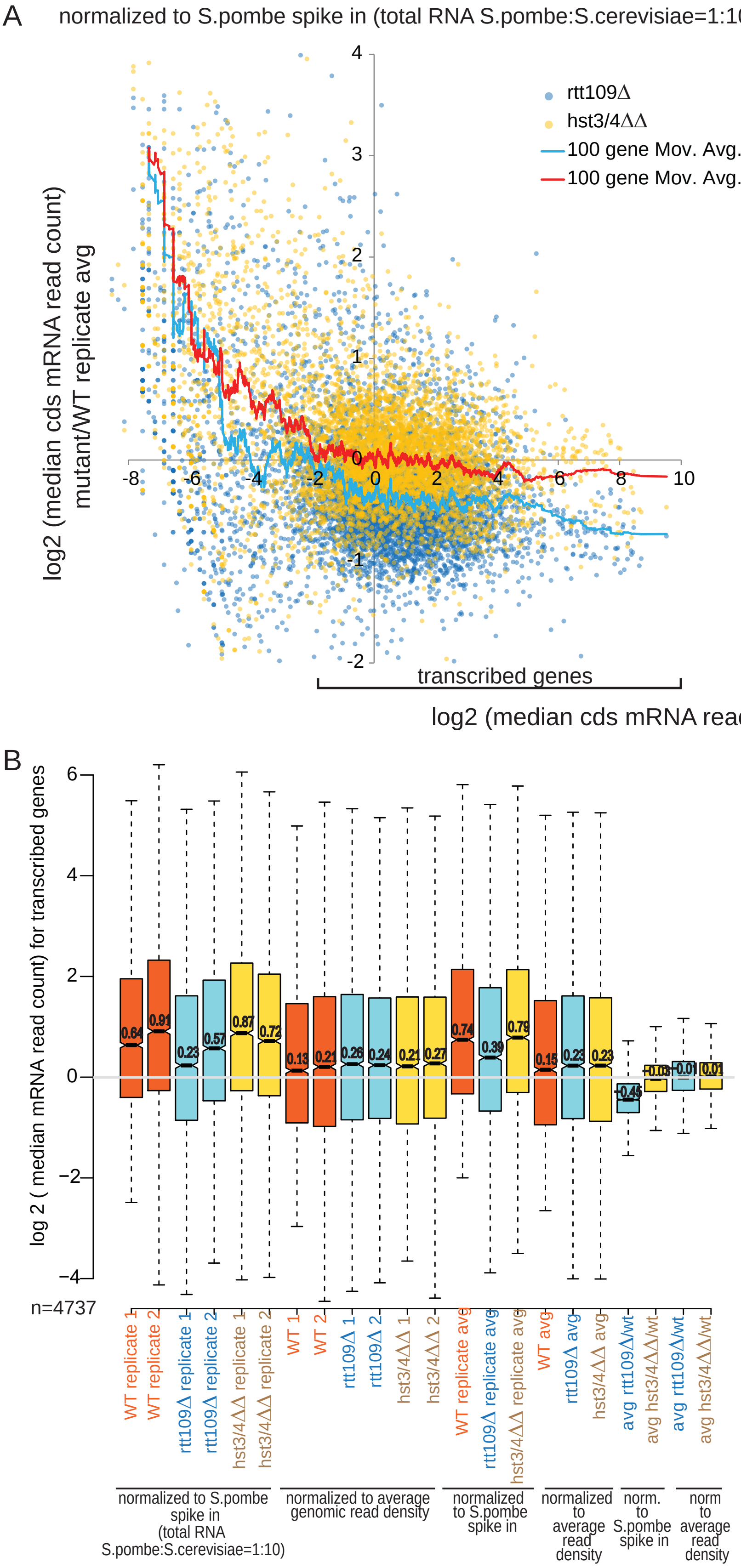


**Figure S5: mRNA levels are globally reduced in the absence of H3K56ac. A.** Total RNA was isolated from wt, rtt109Δ (H3K56 acetylase) and Hst3/4ΔΔ (H3K56 deacetylases) exponentially growing cells (2 replicates each). Total RNA from S.pombe was added to each sample in a 1:10 ratio. Strand specific high-throughput sequencing libraries were then made from isolated polyA RNA. S.Cerevisiae sequencing reads were normalized to the average genomic read density of the S.pombe spike in (left) or of the S.Cerevisiae sample (right). Median sense mRNA levels for each gene (introns were excluded from the calculation) were determined in all samples. The scatter plots show the ratio between mutant and wt mRNA levels for each gene relative to wt mRNA levels. Spike in normalization (left) reveals that mRNA levels in rtt109Δ cells are globally reduced compared to wt, while deletion of hst3 and 4 deactylases has no effect on steady state transcription output. Relative gene expression levels are not affected in either mutant as shown with the internally normalized datasets on the right. **B.** Box plot distributions of median mRNA levels per gene coding region for indicated datasets. Only transcribed genes (shown in A (left panel) were used for the analysis (4737 genes). The average median decrease in global mRNA levels in rtt109Δ cells relative to wt cells is ~30% ($1\text{-}2^{-0.45}$)

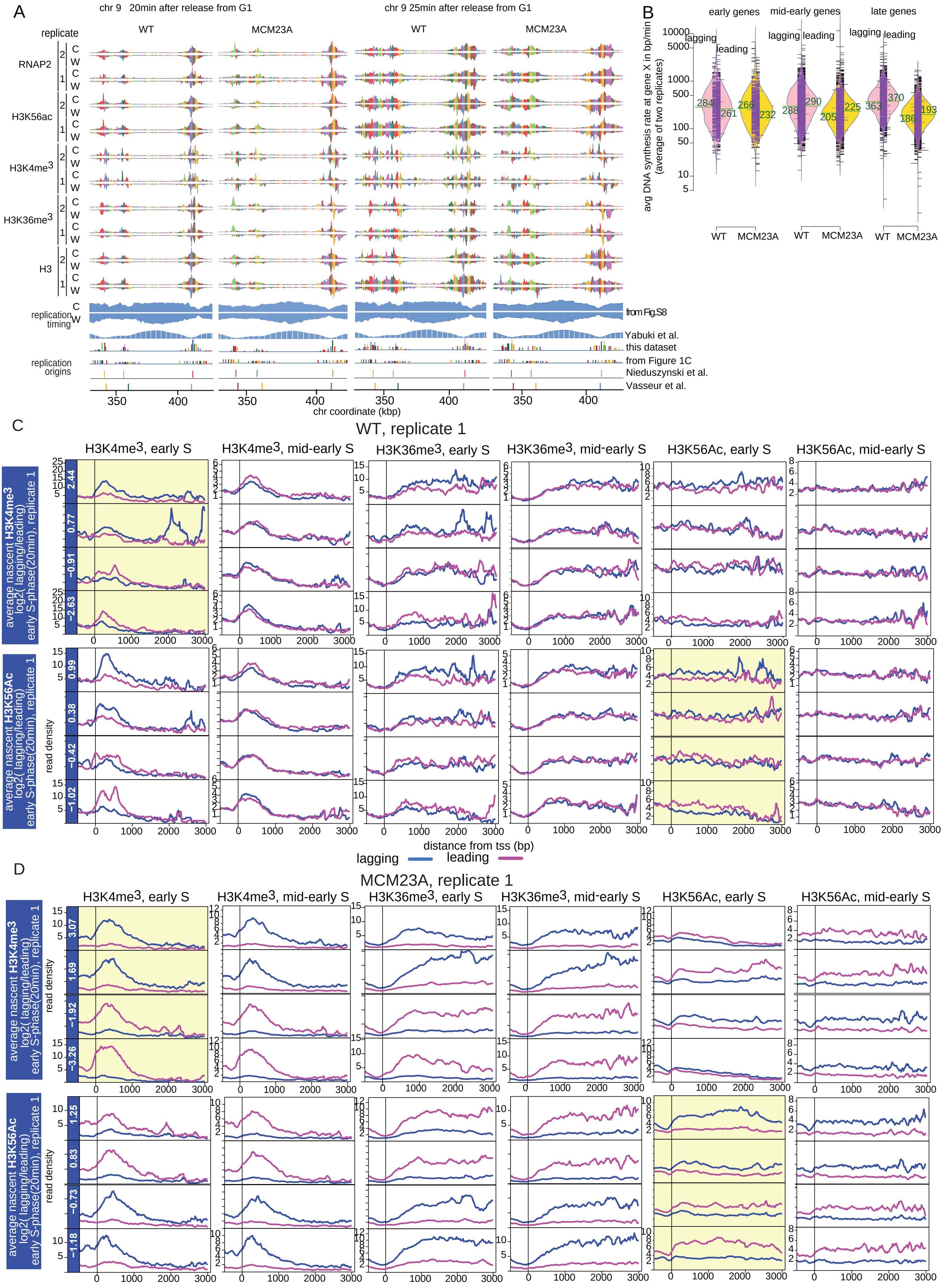
A
chr 9 20min after release from G1
chr 9 25min after release from G1
replicate
WT
MCM23A
RNAP2
H3K56ac
H3K4me3
H3K36me3
H3
C
W
replication timing
from Fig.S8
Yabuki et al.
this dataset
replication origins
from Figure 1C
Nieduszynski et al.
Vasseur et al.
350
400
chr coordinate (kbp)
B
early genes
mid-early genes
late genes
lagging
leading
avg DNA synthesis rate at gene X in bp/min (average of two replicates)
284
261
266
232
288
290
205
225
363
370
186
193
WT
MCM23A
C
WT, replicate 1
H3K4me3, early S
H3K4me3, mid-early S
H3K36me3, early S
H3K36me3, mid-early S
H3K56Ac, early S
H3K56Ac, mid-early S
average nascent H3K4me3 log2( lagging/leading), early S-phase(20min), replicate 1
average nascent H3K56Ac log2( lagging/leading), early S-phase(20min), replicate 1
2.44
0.77
−0.91
−2.63
0.99
0.38
−0.42
−1.02
read density
distance from tss (bp)
lagging
leading
D
MCM23A, replicate 1
3.07
1.69
−1.92
−3.26
1.25
0.83
−0.73
−1.18

**Figure S6 (related to Figures 5, 6 and S7):A.** 400bp binned read density of ChIP-NChAP fractions for RNAP2, H3K56Ac, H3K4me$^{3}$ , and H3K36me$^{3}$ on a segment from chr9 measured in two WT and two MCM23A replicates at 20 (early S) and 25min (mid-early S) after after release from alpha factor G1 arrest. Genes are delimited by different colors. The Watson (W) and Crick(C) strands are counted separately. Read densities were first normalized to the genome wide-average read density for each time-point and then by dividing each segment density with the highest density in the chromosome. Replication timing profiles in the 6th and 7th row from the bottom are taken from the datasets in Figure S8. **B.**Violin plots of Lagging and Leading strand synthesis rates at early, mid-early, and late genes (defined as in Figure 2) for WT and MCM23A strains. The mean for each distribution is shown in green. The black bars represent individual points. Synthesis rates were calculated as described in Figure S8 using Replication Origin Associated Domaines (ROADs) from the 20 and 25min time points from A and replication timing from Figure S8. As in the datasets from figure S8, the bulk of the synthesis rates in MCM23A is slower than in WT. While rates stay constant or even increase for all replication timing gene groups in WT cells they gradually decrease as a function of replication timing in MCM23A mutants. **C.**Average tss centered metagene profiles for WT (replicate 1) ChIP -NChAP fractions indicated on top. Genes from datasets in Fig 5 but only normalized to the genome wide average (i.e. not normalized to input) were divided into 7 bins (only the bottom and the top 2 are shown in the figure) of equal size (~100 genes in each group) according to their average log2(lagging/leading) median read density ratios of early S H3K4me3 (top) or H3K56ac (bottom) ChIP-NChAP fractions from Fig. 5. The value of the average ratio for each bin is indicated in the blue strip on the left. **D.** Same as C but for MCM23A (dataset from Fig. 6).

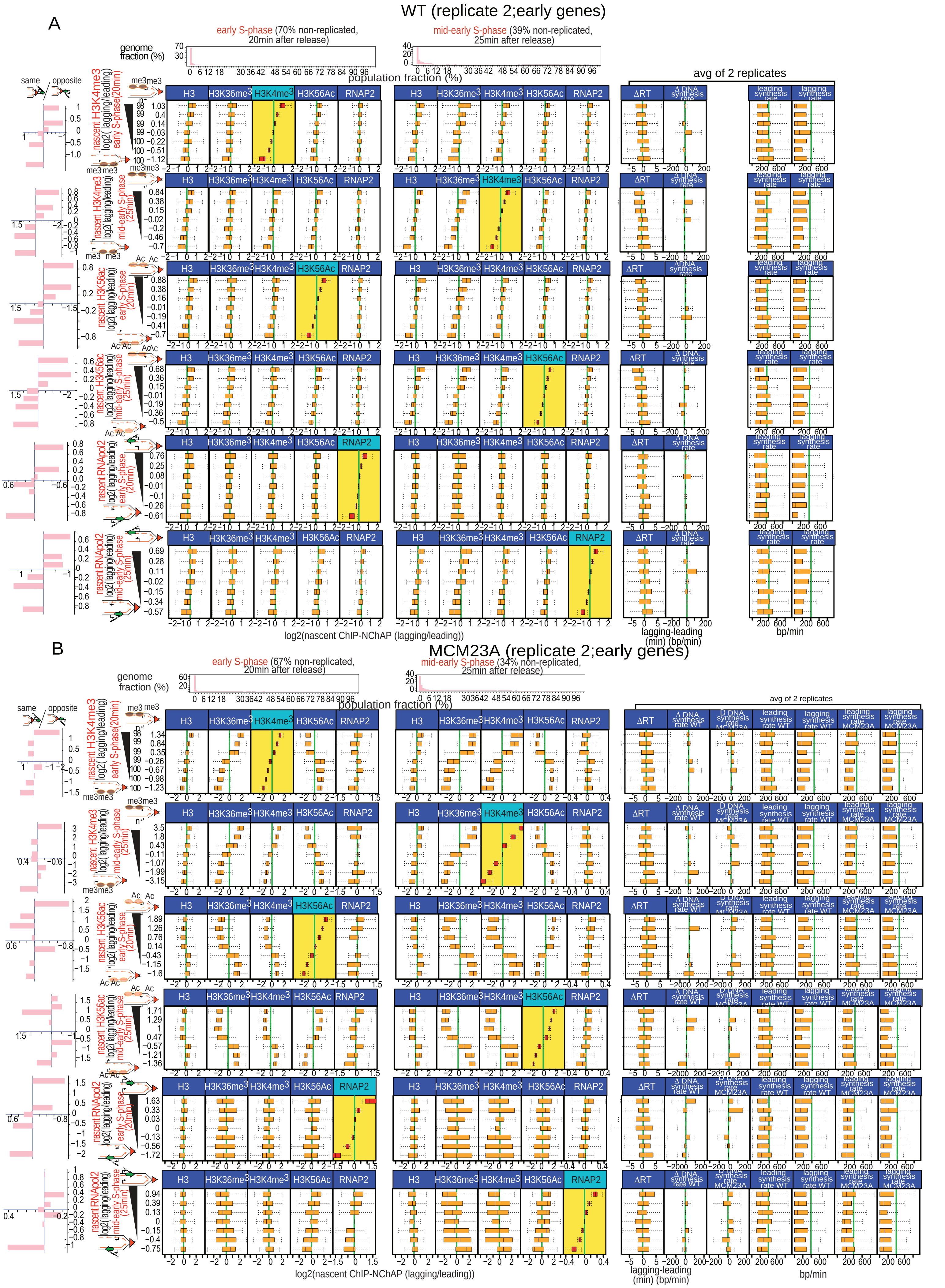


**Figure S7: Box plot distribution of H3, H3K4me3, H3K36me3, H3K56ac and RNApol2 on the lagging and leading strand.** Biological replicate 2 for wt (A) and MCM23A (B) mutant cells. The lay out is the same as in Figures 5 and 6.

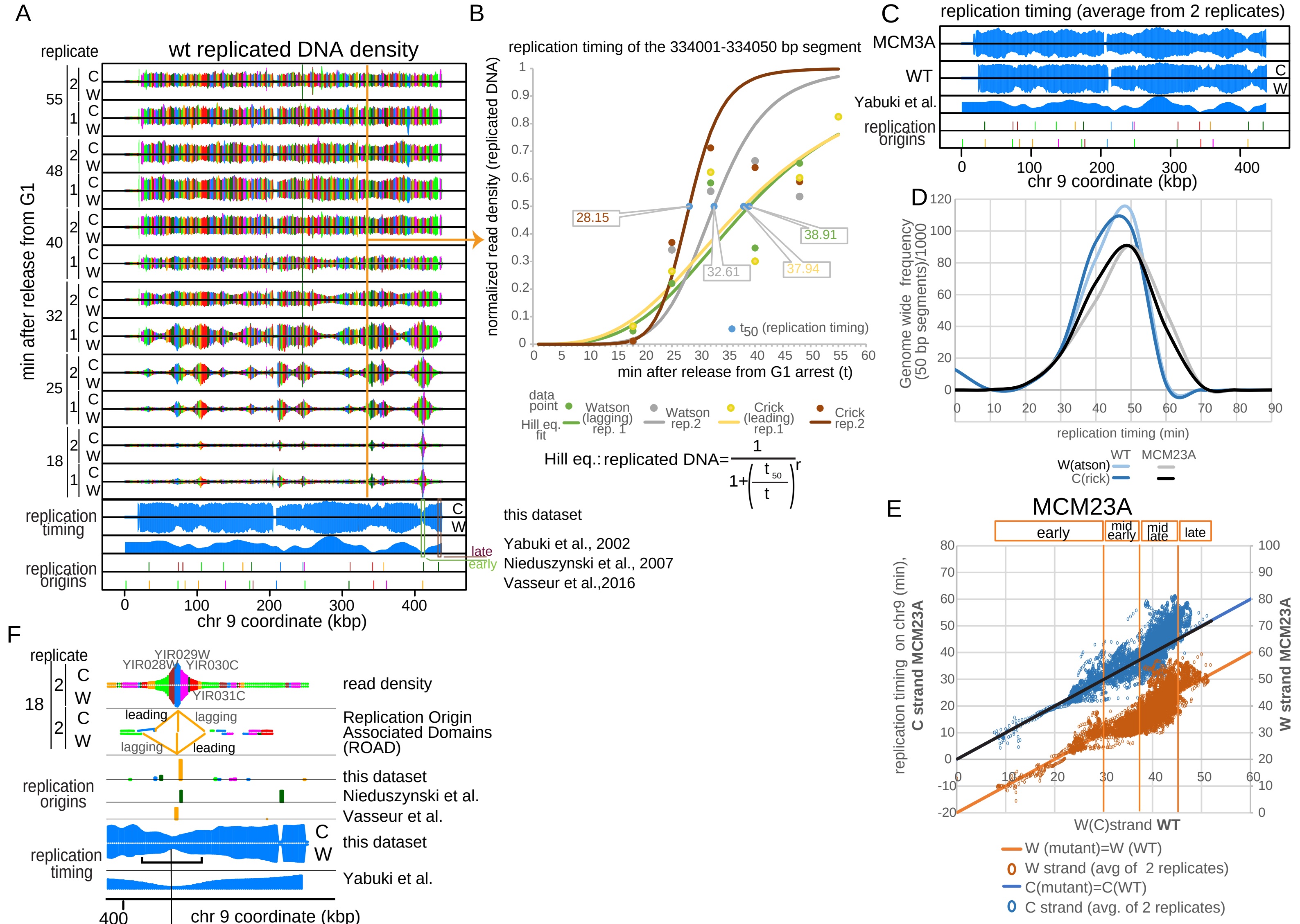


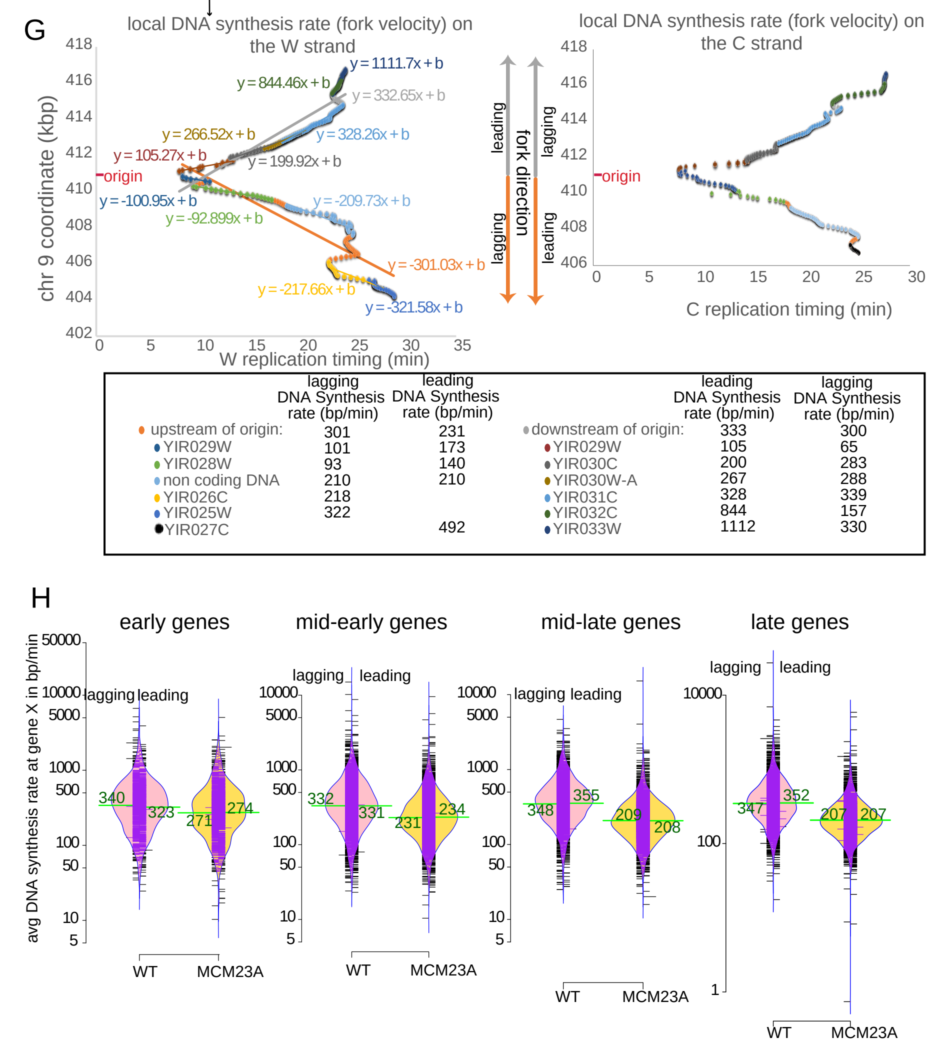


**Figure S8: Calculations of replication timing and DNA synthesis rates per gene for WT and MCM23A strains. A.** 400bp binned input normalized read density of replicated DNA (NChAP) for chr9 measured in two WT replicates from a timecourse after release from alpha factor G1 arrest. Genes are delimited by different colors. The Watson (W) and Crick(C) strands are computed separately. To calculate replication timing shown in the 4th and 5th rows from the bottom reads from each time point were binned in 50bp segments and normalized by dividing each segment density with the highest density in the chromosome. The normalized densities were then smoothed with a 1600bp moving window average. Read densities for each 50bp segment were then plotted against time as shown in B. **B.** Replication timing $t_{50}$ from the Hill equation fit to the time course data points as shown.**C.** Replication timing profiles of chr9 for Watson and Crick strands of WT and MCM23A strains. **D.** Replication timing distribution of all 50bp segments in the genome for WT and MCM23A. **E.** MCM23A replication timing vresus WT replication timing for the Watson strand (right Y axis) and the Crick strand (left Y axis). All late replicating and at a portion of mid-late replicating genes replicate ~10min later in the MCM23A mutant compared to wt. **F-G.** Replication timing values were used to calculate DNA synthesis rates of the Watson and Crick strands at every replicated gene from the 18 and 25min time points from A. **F.** First replication origins were determined from local peaks in the 18 et 25min time points. Second, genes that are replicated from any given origin were grouped into Replication Origin Associated Domains (ROADs) for either time point as illustrated **G.**The average DNA synthesis rates of the Watson (left) and Crick (right) strands at each gene from every ROAD were calculated from the slope of the linear fit of the gene coordinates in bp versus replication timing as shown in the graphs Synthesis rates of lagging strand gene copies are derived from slopes from the Watson strand upstream of origins and the Crick strand downstream of origins. Conversely leading strand synthesis rates come from Crick strands upstream and Watson strands downstream of origins.Synthesis rates for genes associated with the ROAD from F are summarized in the table below the graphs.**H.** Violin plot of Lagging and Leading strand synthesis rate distribution at early, mid-early, mid-late and late genes (defined as in E) for WT and MCM23A strains. The mean for each distribution is shown in green. The black bars represent individual data points. The bulk of synthesis rates in MCM23A is lower than in WT, and while rates stay constant for all replication timing gene groups in WT cells they gradually decrease as a function of replication timing in MCM23A mutants.

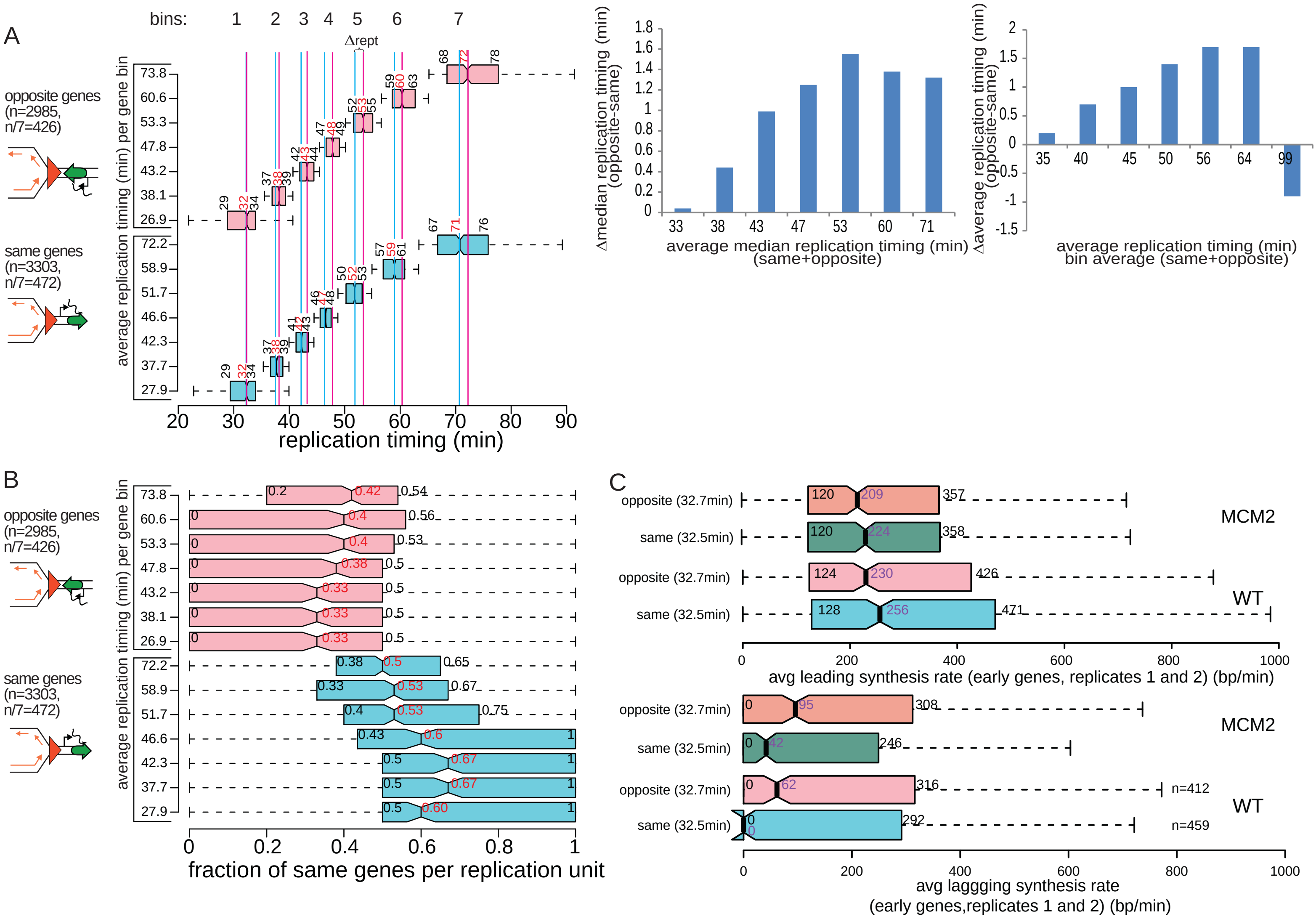


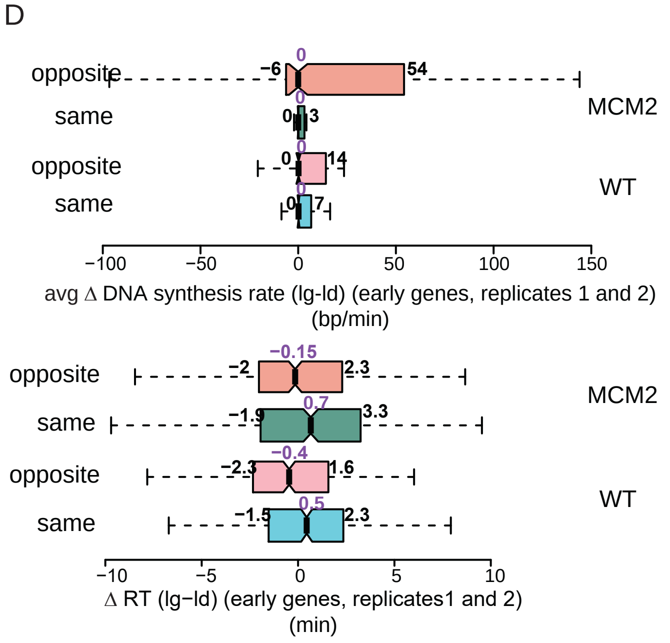


**Figure S9: A.** Replication timing differences of "same" and "opposite" genes 6288 yeast genes were divided into "same" (3303 genes) and "opposite" (2985 genes) orientation genes, sorted by replication timing (Vasseur et al., 2016) and then divided into seven equal bins each (numbered above the plot, 426 and 472 genes per bin for opposite and same genes, respectively, left panel). Left panel: box plot distributions of replication timing were determined for each bin. Second and third quartile medians are shown in black above each box. Medians for the entire distribution are shown in red. Δrept (marked on bin 5) is the difference in median replication timing between "opposite" (pink) and "same" (blue) genes. Middle and right panels: Bar graph of the differences in median (middle panel,red values in left panel) or average replication timing (right panel) between opposite and same genes for each bin. The x axis shows the average of medians (middle) or average replication timing (right) of same and opposite gene bins (1 to 7). **B.** Genic orientation bias of replication units.The ensemble of all the genes that replicated from the same closest replication origin and are on the same side of that origin (upstream or downstream) define each replication unit. To determine whether any given gene is more likely to be surrounded by genes of the same genic orientation within each replication unit, the fraction of "same" orientation genes was calculated for each replication unit. We then determined box plot distributions of "same" gene fractions from replication units assigned to genes from each replication timing bin defined in A. Second and third quartile medians are shown in black and medians for the entire distribution are shown in red. **C.** Box plot distribution of lagging (bottom) and leading (top) strand DNA synthesis rates for "same" (n=459) and "opposite" (n=412) early replicating genes in wt and mcm23A mutants. The average replication timing for the gene group (from Vasseur et al., 2016) is in parenthesis on the left. Synthesis rates are an average of two replicates (replicates 1 from Figures 5 and 6 and replicates 2 from Figure S8). The median values are shown in purple and the 2nd and 3rd quartiles in black. **D.** Top: Box plot distribution of the difference in DNA synthesis rates between lagging and leading (lg-ld) copies of "same" and "opposite" early genes as in C. Bottom: Distribution of the differences between replication timing (RT) of lagging and leading strands from the same group of genes as in C.

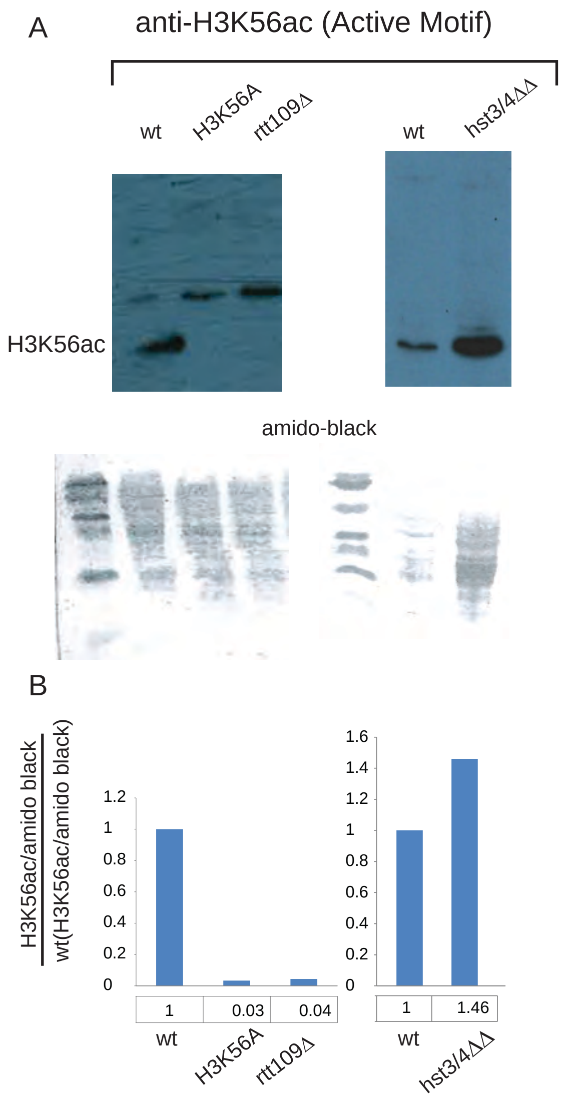


**Figure S10: Specificity test for the anti H3K56ac antibody (Active Motif). A.** Western blots with midlog cell extracts from indicated strains. wt: w303 (MATa ade2-1 his3-11,15 leu2-3,112 trp1-1 ura3-1 can1-100); rtt109Δ: strain RZ72; H3K56A: AC33; hst3/4ΔΔ: AC29; Genotypes are listed in Table S3.The bottom panel shows the amido-black staining of the gels in the top panel.**B.** Quantification fo western blots in A. H3K56ac background corrected band intensities were divided by the intensity of the corresponding lane stained with amido black. Band intenisties were determined using the Gel Analyzer software (GelAnalyzer.com, Istvan Lazar). The background signal was determined with the rolling ball method (radius= 50).
